# Evidence for conservation of primordial ∼12-hour ultradian gene programs in humans under free-living conditions

**DOI:** 10.1101/2023.05.02.539021

**Authors:** Bokai Zhu, Silvia Liu, Natalie L. David, William Dion, Nandini K Doshi, Lauren B. Siegel, Tânia Amorim, Rosemary E. Andrews, GV Naveen Kumar, Hanwen Li, Saad Irfan, Tristan Pesaresi, Ankit X. Sharma, Michelle Sun, Pouneh K. Fazeli, Matthew L. Steinhauser

**Affiliations:** Aging Institute of UPMC, University of Pittsburgh School of Medicine; Pittsburgh, Pennsylvania, USA; Pittsburgh Liver Research Center, University of Pittsburgh; Pittsburgh, Pennsylvania, USA; Division of Endocrinology and Metabolism, Department of Medicine, University of Pittsburgh School of Medicine; Pittsburgh, Pennsylvania, USA; Department of Pathology, University of Pittsburgh School of Medicine; Pittsburgh, Pennsylvania, USA; Neuroendocrinology Unit, Division of Endocrinology and Metabolism, Department of Medicine, University of Pittsburgh School of Medicine; Pittsburgh, Pennsylvania, USA; Center for Human Integrative Physiology, University of Pittsburgh School of Medicine; Pittsburgh, Pennsylvania, USA; Department of Statistics, Kenneth P. Dietrich School of Arts and Sciences, University of Pittsburgh; Pittsburgh, Pennsylvania, USA; Division of Cardiology, Department of Medicine, University of Pittsburgh School of Medicine; Pittsburgh, Pennsylvania, USA

## Abstract

While circadian rhythms are entrained to the once daily light-dark cycle of the sun, many marine organisms exhibit ∼12h ultradian rhythms corresponding to the twice daily movement of the tides. Although human ancestors emerged from circatidal environment millions of years ago, direct evidence of ∼12h ultradian rhythms in humans is lacking. Here, we performed prospective, temporal transcriptome profiling of peripheral white blood cells and identified robust ∼12h transcriptional rhythms from three healthy participants. Pathway analysis implicated ∼12h rhythms in RNA and protein metabolism, with strong homology to the circatidal gene programs previously identified in Cnidarian marine species. We further observed ∼12h rhythms of intron retention events of genes involved in MHC class I antigen presentation, synchronized to expression of mRNA splicing genes in all three participants. Gene regulatory network inference revealed XBP1, and GABP and KLF transcription factor family members as potential transcriptional regulators of human ∼12h rhythms. These results suggest that human ∼12h biological rhythms have a primordial evolutionary origin with important implications for human health and disease.

## Introduction

Biological rhythms are conserved from single-cell organisms to humans. The relevance of biological rhythms is exemplified by circadian rhythms, which are entrained to a ∼24-hour cycle by the rotation of the Earth and daily exposure to sunlight. Circadian gene expression programs in turn regulate canonical hormonal systems, vasomotor activity, coagulation, amongst other processes, accounting for predilection of certain disease events at specific times of day [1]. Genetic variants in core circadian genes and disruptions of circadian rhythms are associated with increased risk of diverse human diseases [1–4]. The wide-ranging pathological effects of circadian disruption demonstrate the centrality of the 24h circadian clock to homeostasis and disease pathobiology [1].

Our understanding of biorhythms is expanded by alternative, non-circadian rhythms in lower organisms. Coastal marine organisms, such as the sea anemone *A. diaphana,* exhibit ∼12h ultradian rhythms, entrained by the twice daily movements of the tides [5]. In mice, many ∼12h rhythms of gene expression involved in mRNA, protein and lipid metabolism are established by an XBP1-dependent oscillator independent of the circadian clock or cell cycle [6, 7]. Oscillation of some physiological metrics at an approximate ∼12h interval in humans suggests the possibility of a corresponding 12h molecular oscillator [6]. However, direct evidence for ∼12h rhythms at the molecular level in humans is lacking. In a recent single time-point examination of postmortem human brain tissues using time-of-death as surrogate for circadian time, we uncovered ∼12h ultradian rhythms of gene expression in dorsolateral prefrontal cortex regions that are implicated in unfolded protein response (UPR) and mitochondria-related metabolism like oxidative phosphorylation [8]. While this study provided support for the existence of ∼12h rhythms in humans, time-of-death may not accurately reflect internal circadian time. To overcome this limitation, we performed a prospective longitudinal study of three healthy participants admitted to an inpatient clinical research unit for high temporal resolution (q2h) sampling of peripheral white blood cells. We discovered a shared ∼12h gene program with homology to our ancient marine ancestors and involving fundamental pathways of protein and RNA metabolism.

## Results

### Gene expression oscillations in human white blood cells under free-living conditions

We studied 3 healthy males (**Table 1**) who self-reported a regular nighttime sleep schedule and did not engage in nighttime shift work or other sleep-disrupting activities and who had a body mass index between 18.5-24.9 kg/m^2^. Participants were admitted for a 1-day acclimatization period prior to 48 hours of blood sampling at a 2-hour sampling interval (**Fig. 1A**), a frequency that allowed detection of oscillations with periods of 6 hours or longer with high confidence [9, 10]. The protocol reflected our aim to sample with sufficient frequency and duration to detect ∼12h rhythms within individuals. We also sought to mimic free-living conditions by feeding them a diet similar in caloric content and macronutrient composition to their typical intake and encouraging maintenance of their routine sleep/wake cycles. Overnight, blood was collected via a long intravenous line from outside the room to avoid exposing participants to light or sleep disruption.

**Table 1.** Participant characteristics.

|  | Participant 1 | Participant 2 | Participant 3 |
| --- | --- | --- | --- |
| Age (years) | 20 | 22 | 28 |
| Body mass index (kg/m<sup>2</sup>) | 23.3 | 23.7 | 24.4 |
| Vitamin D (ng/mL) | 85 | 26 | 22 |
| Glucose (mg/dL) | 73 | 84 | 74 |
| BUN (mg/dL) | 10 | 18 | 14 |
| Creatinine (mg/dL) | 0.9 | 1.0 | 0.9 |
| AST (U/L) | 22 | 41 | 22 |
| ALT (U/L) | 10 | 28 | 22 |
| TSH (mIU/L) | 1.306 | 1.48 | 0.803 |

**Fig. 1.**
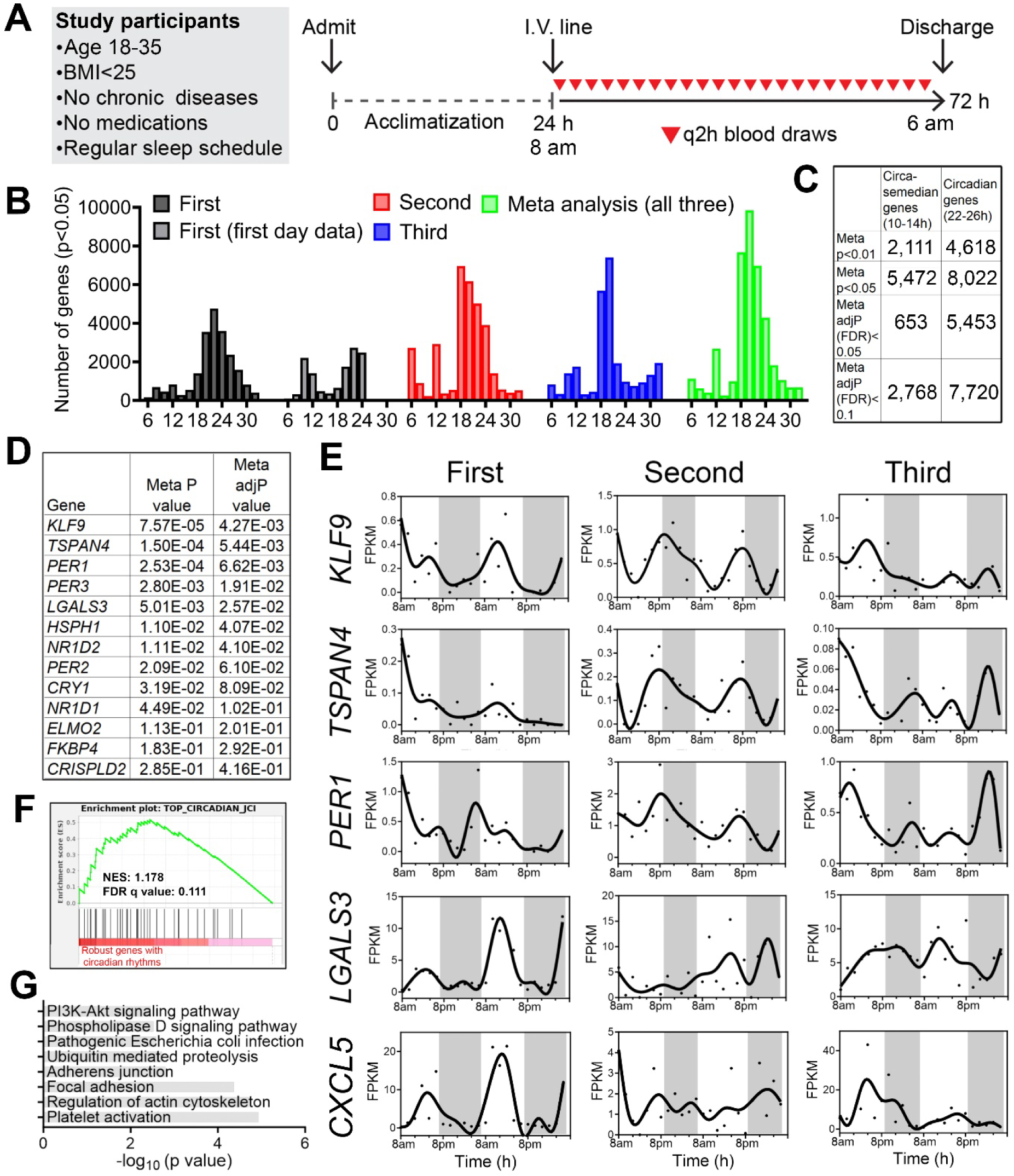
Identification of inter-individual variability of circadian and ultradian rhythms of gene expression in peripheral white blood cells of humans. (**A**) Human protocol and study schematic. (**B-D**) Histograms showing the period distributions of all rhythmic genes uncovered from the three participants (with p value<0.05) as well as the period distribution of rhythmic genes calculated by the meta p values (meta p<0.05). For the first individual, periods calculated from all 48 hours of data or the first 24h hours of the dataset were both shown. (**C**) Table summarizing the number of circadian and ∼12h genes with different statistical cut-offs. (**D**) Table listing the meta P and meta adj-P (FDR) values of thirteen circadian genes previously used as training dataset for predicting circadian phase from human blood samples [14]. (**E**) Raw temporal expression profile (dot) and spline fit (solid line) of five of the thirteen circadian genes. (**F**) Gene set enrichment analysis (GSEA) showing enrichment of a previously identified set of top forty robustly-cycling circadian genes in human white blood cells [14] with robust circadian rhythmicity in our study. (**G**) GO analyses of top KEGG pathways for circadian genes with meta adj-P (FDR) less than 0.01.

We performed bulk mRNA-Seq in buffy coat fractions prospectively collected at 2h intervals for 48 hours (24 samples/participant) (**Fig. 1A**) (**Table S1**). To identify genes cycling with a period ranging from 6 to 32 hours, we applied the RAIN algorithm to each individual’s temporal transcriptome [11]. Compared to other competing methods like JTK_CYCLE, RAIN detects rhythms with arbitrary waveforms and therefore more robustly uncovers ultradian rhythms [10, 12, 13]. As shown in **Fig. 1B**, we observed inter-individual variability in the number of genes cycling at different periods. Specifically, the first individual exhibited dominant oscillations cycling at the circadian period between 20-24 hours, with a secondary population detected at the periods of 10-12 hours. These 10-12h oscillations were more evident in the first 24 hours, with dampening of ultradian rhythms on the second day (**Fig. 1B**). Aside from circadian oscillations between 18 and 24 hours, the second participant exhibited ultradian rhythms cycling at periods of 6 and 12-hours (**Fig. 1B**). The third individual also exhibited oscillations cycling at 6-hours, 12-hours, and at ∼20 hours, with very few genes observed cycling at 24-hours (**Fig. 1B**). The inter-individual variability of the oscillating transcriptome profiles in our study may be attributable to the fact that the participants were held at living conditions that resemble free-living. By contrast, in a prior study that prospectively examined circadian gene expression in human white blood cells, participates were maintained in a semi-recumbent position under dim light and constant temperature and humidity during 40 hours of sleep deprivation [14].

To increase statistical power to detect oscillations common to all three individuals, we performed a meta-analysis and generated a combined p-value for each gene at periods from 6- to 32-hours using Fisher’s method. This method has been extensively used in medical and genetic research to combine the results from independent tests with the same null hypothesis [15]. In our case, each participant was distinct and studied at a different time, however the null hypothesis was the same: the absence of rhythms. Using combined p-values (herein referred to as meta p values) with an alpha of 0.05, we observed two major populations of oscillations cycling at 12- and 18-24-hours, with the largest number of genes observed cycling at 20-hours (**Fig. 1B**). Using the Benjamini-Hochberg procedure to further adjust for the false discovery rate on meta p values (meta adj-P value or FDR), we uncovered 653 circasemidian (∼12h) and 5,453 circadian (∼24h) genes with an FDR less than 0.05 (**Fig. 1C**) (**Table S2**).

### Human white blood cells exhibit rhythmicity in canonical circadian genes

To determine the soundness of our RNA-seq dataset, we compared our putative circadian gene set to a prior study, where participants were maintained in much more stringently controlled conditions of dim-lighting, fixed semi-recumbent positioning, and sleep deprivation [14]. In that study, a total of thirteen circadian genes exhibiting robust circadian expression in peripheral white blood cells were utilized as a training gene set to predict the circadian phases from single time point blood samples [14], of which seven and nine exhibited meta adj-P values less than 0.05 and 0.1, respectively, in our study (**Fig. 1D**). This set includes canonical circadian clock genes *PER1*, *PER2*, *PER3*, *NR1D1*, *NR1D2* and *CRY1* (**Fig. 1D, E**). Gene set enrichment analysis (GSEA) further indicated that the forty circadian genes that cycled in at least 50% of the human participants in that study also exhibited circadian oscillations with low meta p values in our study (**Fig. 1F**). For these core circadian genes we observed inter-individual variability in the phases and amplitudes among the three participants (**Fig. 1E**), likely reflecting individual-specific sleep-wake schedules. Interestingly, even under constant routine conditions, these genes also exhibited inter-individual variability in phase and amplitudes, albeit to a much less degree (**Fig. S1**). Overall, circadian genes in human peripheral white blood cells are strongly involved in pathways of platelet activation, regulation of actin cytoskeleton and adhesions (**Fig. 1G**). Collectively, these results demonstrated the robustness of our study design and dataset, further justifying the treatment of each longitudinal time series as an independent dataset followed by a meta-analysis to achieve greater statistical power.

### ∼12h ultradian rhythms of gene expression are involved in mRNA and protein metabolism in human peripheral white blood cells

Oscillations cycling close to a 12-hour period were the second most abundant in all three individuals after the circadian population (**Fig. 1B**). Using meta adj-P values (FDR) as cut-off, we identified 653 and 2,768 ∼12h genes (with period between 10-14-hours) with meta adj-P (FDR) less than 0.05 and 0.1, respectively (**Figs. 1C and 2A**). Importantly, the average gene expression profiles of these 653 (FDR<0.05) and 2,115 (0.05<FDR<0.1) genes are very similar to each other in each of the three individuals (**Fig. 2A**).

**Fig. 2.**
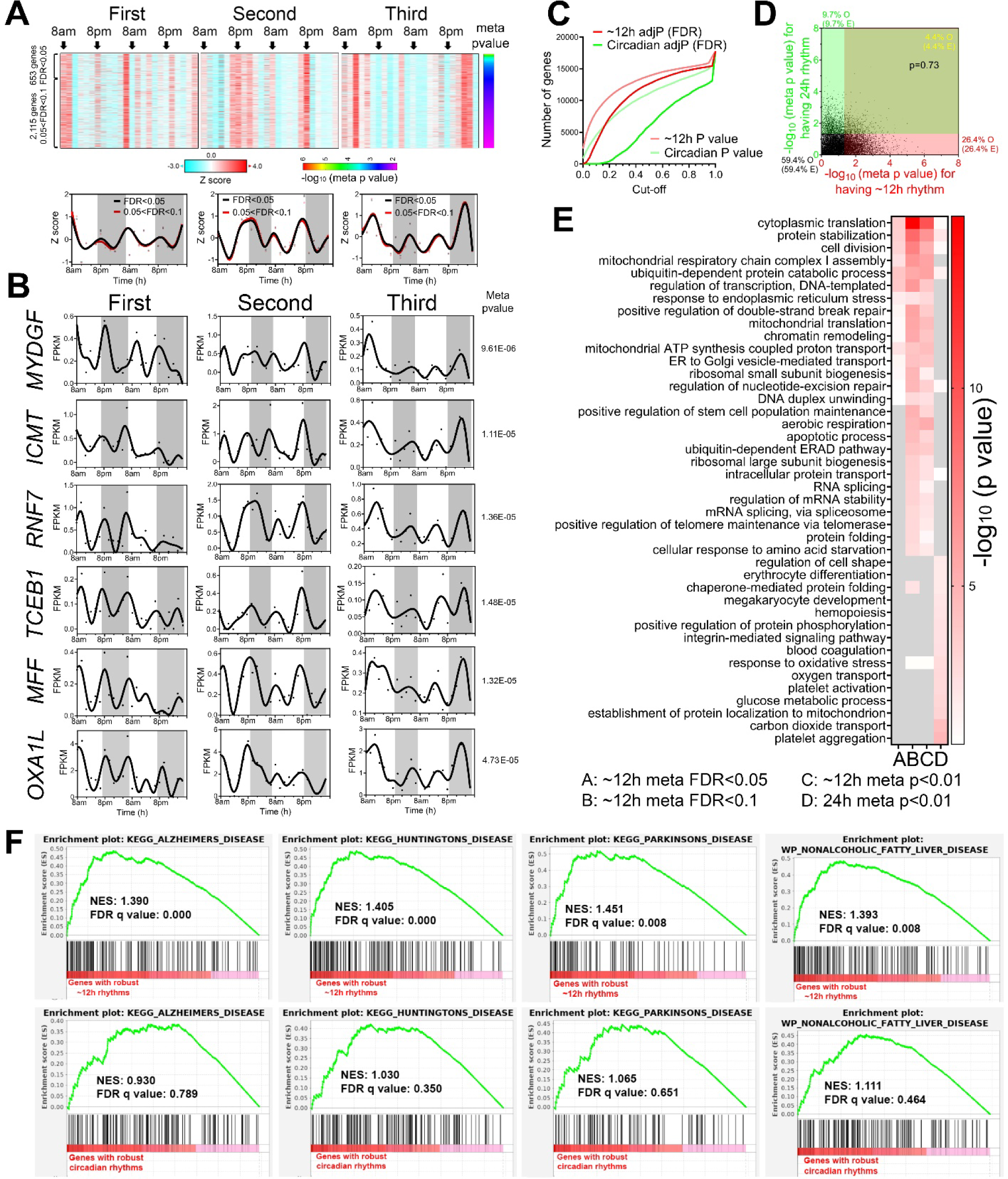
∼12h gene expression rhythms are enriched in mRNA and protein homeostasis pathways. (**A**) Heatmap and quantification of ∼12h genes from all three participants with meta adj-P (FDR) values less than 0.05 or between 0.05 and 0.1. (**B**) Raw temporal expression (dot) profile and spline fit (solid line) of six ∼12h genes with meta p values also shown. (**C**) Cumulative distribution of the number of circadian and ∼12h genes with different meta p and meta adj-P cut-offs. The circadian data set consisted of a 4h sampling interval so that the period to sampling interval was matched to the ∼12h data set. (**D**) Scatter plot comparing log transformed meta p values for each gene exhibiting circadian versus ∼12h rhythms. Both observed and predicted percentage of genes (under the null hypothesis that ∼12h and circadian genes are independently regulated) with meta p value smaller or larger than 0.05 are further shown. P value of 0.73 is calculated by Chi-square test. (**E**) GO analysis of circadian and ∼12h genes with different statistical cut-offs. (**F**) GSEA showing enrichment scores of different gene sets for ∼12h (top) and circadian (bottom) genes.

As observed with circadian genes, inter-individual variability was also observed for ∼12h ultradian rhythms. In the first and second participants, the acrophases (time of gene expression peak) were at 8 am and 8 pm, while ∼12h rhythms in the third individual peaking around 4 am and 4 pm (**Fig. 2A**). Examples of robust ∼12h genes cycling in all three individuals include *MYDGF*, *ICMT*, *RNF7*, *TCEB1*, *MFF* and *OXA1L* (**Fig. 2B**). *MYDGF* encodes an endoplasmic reticulum (ER)-localized protein with secreted forms also identified from monocytes/macrophages that promotes tissue repair in a murine model of myocardial infarction [16, 17]. *ICMT* encodes a protein-*S*-isoprenylcysteine *O*-methyltransferase that also localizes in the ER and is responsible for cell membrane-targeting of selective proteins [18]. *RNF7* encodes an essential subunit of SKP1-cullin/CDC53-F box protein ubiquitin ligases [19] and TCEB1 protein is a subunit of the transcription factor B responsible for transcription elongation [20]. Both MFF and OXA1L are mitochondrial proteins localized to outer and inner mitochondria membranes, respectively, and are essential for mitochondria fission and assembly of respiratory complex I [21, 22].

One important question regarding ultradian rhythmicity is whether it reflects ‘real’ rhythms with biological significance, or simply mathematical artefact. This is particularly relevant for rhythms that cycle at harmonic frequencies of the 24-hour circadian period, as non-sinusoidal circadian waveforms (seesaw-like or square-shaped waveforms) often have superimposed ∼12 and ∼8-hour harmonic frequencies detected by spectral analytical methods such as a Fourier transformation, eigenvalue/pencil, or wavelet analyses [12, 23–25]. We took several approaches to address this question. First, 12-hour transcriptional oscillations did not cycle at the exact second harmonic of the dominant circadian period in any of the three participates: the first harmonic circadian periods were 22, 18 and 20-hours in the three individuals, respectively, and 20-hours for the meta-analysis (**Fig. 1B**). Second, if ∼12h oscillations are indeed harmonics of the circadian rhythm, then genes exhibiting both circadian and ∼12h ultradian rhythmicity should be detected at a higher frequency than expected by chance. To test this, we generated a circadian gene set using a 4-hour interval subset of the original dataset (8am, 12am, 4pm, etc.) to achieve an equivalent number of data points per period—i.e. equivalent sensitivity—and a meta-p cut-off of 0.05. Importantly, the circadian subset showed a strong positive correlation with meta-p values obtained with the full 2h interval dataset (**Fig. S2A**). After this adjustment to match sampling interval to period ratios, ∼12h gene rhythms were more prevalent than circadian genes (**Fig. 2C**). Moreover, the expected and observed percentages of genes having circadian and/or ∼12h rhythms were nearly identical (p=0.73 by Chi-square test), indicating independent detection of ∼12h and circadian rhythms (**Fig. 2D**). Third, we performed Gene Ontology (GO) and GSEA to compare biological pathways enriched in circadian and ∼12h genes, respectively. We reasoned that if ∼12h oscillations are mathematical harmonics of the circadian rhythm, then ∼12h and circadian genes should share enriched biological pathways. Both analyses revealed largely distinct pathways associated with circadian and ∼12h rhythms (**Figs. 2E** **and** **S2B**). While circadian rhythms were enriched in platelet activation, blood coagulation and cell adhesion as previously demonstrated (**Figs. 2E** **and S2B**), ∼12h genes were associated with fundamental biological processes of mRNA (such as spliceosome and RNA splicing), protein metabolism (including ubiquitin-mediated proteolysis, response to ER stress, protein folding and protein transport in the ER and Golgi), and mitochondria complex chain assembly (**Figs. 2E** **and S2B**). Cytoplasmic translation, on the other hand, was enriched in both circadian and ∼12h genes (**Fig. S2B**). Consistent with the known causal roles of dysregulated proteostasis in proteinopathies, and impaired mitochondrial metabolism in metabolic syndromes, ∼12h genes were also much more strongly enriched in neurogenerative disease gene sets and nonalcoholic fatty liver disease (**Fig. 2F**). Enrichment of protein metabolism and mitochondrial respiration genes is also in alignment with ∼12h gene signatures in human dorsolateral prefrontal cortex [8].

We next performed pathway analyses of the ∼12h gene sets for each individual participant. We either analyzed the data as a continuous 48h time series with a single data point at each time point (RAIN conti) or as a 24h time series where data points collected at the same time on two consecutive days were treated as biological replicates (RAIN dupli) (**Table S2**). The RAIN duplicate approach revealed larger ∼12h gene programs (p value cut-off of 0.05) with lower FDRs: 3,462 genes (FDR=0.224), 7,060 genes (FDR=0.119) and 4,807 genes (FDR=0.166) (**Figs. S3 and 4**). As expected, ∼12h genes common to all three individuals also tended to have small meta-p values (**Fig. S3E and Fig. S4E**). Pathways related to mRNA and protein metabolism emerged as significantly enriched for 12h rhythm genes in each participant regardless of the inputs or thresholds for RAIN analysis (**Figs. S3G and S4C, E**).

To determine robustness to different analytic methods, we also performed spectrum analysis with the eigenvalue/pencil method [10, 12, 13, 26–28], which unlike statistical methods such as JTK_CYCLE and RAIN does not require pre-assignment of period range and thus allows unbiased identification of multiple oscillations for any given gene [10, 12, 13, 26–28]. Eigenvalue/pencil analyses also revealed prevalent circadian and ∼12h oscillations in all three individuals (**Figs. S5-7**, **and Table S3**). Since the eigenvalue/pencil algorithm involves non-statistical signal processing, we used a permutation-based method that randomly shuffles the time label of gene expression data to determine the false discovery rate (FDR) for the 12h gene list [13, 29]. The FDR for the ∼12h gene lists was estimated to be 0.33, 0.36 and 0.38 for each of the three participants respectively (**Figs. S5F, S6F, and S7F**). It is important to note that the calculated FDR is likely an overestimation since amplitude and phase information for the ∼12h rhythms is not considered. Application of eigenvalue/pencil analyses confirmed that 1) ∼12h gene expression was dampened in the second day in the first individual but remained steady over 48 hours in the second and third participants (**Figs. S5G, S6G, and S7G**), 2) ∼12h rhythms peaked around 8am and 8pm in the first and second individuals, and around 4am and 4pm in the third (**Figs. S5C, S6C and S7C**), and 3) ∼12h genes were strongly enriched in mRNA and protein metabolism pathways and distinct from the circadian gene sets in all three individuals, (**Figs. S5J, S6J and S7J**). In addition, we found that the relative amplitude (amplitude normalized to the mean expression of each gene) of ∼12h rhythms in all three individuals are less than their respective circadian counterparts, with mean fold change of 2.74, 1.88 and 1.67 observed in the three individuals (**Figs S5E, S6E, and S7E)**. The smaller relative amplitude of ultradian compared to circadian gene expression rhythms has been also observed in mice [12], and may be due to physical restraints on the maximal rates of RNA polymerase II-mediated transcription can achieve [12]. Taken together, multiple analytical methods and statistical thresholds provided convergent evidence supporting the existence of ∼12h rhythms of gene expression implicated in mRNA and protein metabolism in human white blood cells.

### Identification of candidate transcriptional regulators of ∼12h rhythms

To infer gene regulatory networks governing ∼12h rhythms in human white blood cells, we performed LISA [30] and motif analysis on the top 500 ∼12h genes with the lowest meta-p values. We cross-referenced enriched motifs with transcription factor genes that exhibited ∼12h rhythms (with meta-adjP<0.1), identifying XBP1, CREBZF, GABPB1/2, NFYB, GMEB1, KLF16 as top candidates that are preferentially enriched with ∼12h genes compared to circadian genes (**Fig. 3A-D**). XBP1, CREBZF and CREB1 belong to the Basic Leucine Zipper (bZIP) transcription factor family and are established transcription regulators of the UPR and adaptive stress response [31–35]. GABPB1/B2 are ETS family transcription factors involved in mitochondria biogenesis and oxidative phosphorylation [36, 37]. NFYB encodes one of the three subunits of nuclear transcription factor Y (the other being NFYA and NFYC) and its DNA binding motif was recently shown to co-occur with XBP1 in mouse type-2 T helper cells [38]. Importantly, that many of these transcription factors have been implicated in regulation of ∼12h rhythms in mice. For instance, it was previously proposed that ∼12h rhythms are generally established by a tripartite network comprising ETS, bZIP and NFY transcription factors, whereas KLF transcription factors may control ∼12h rhythms in a more tissue specific manner [39]. Enriched GMEB DNA binding motifs were also observed in the promoters of hepatic ∼12h rhythms in mice [26].

**Fig. 3.**
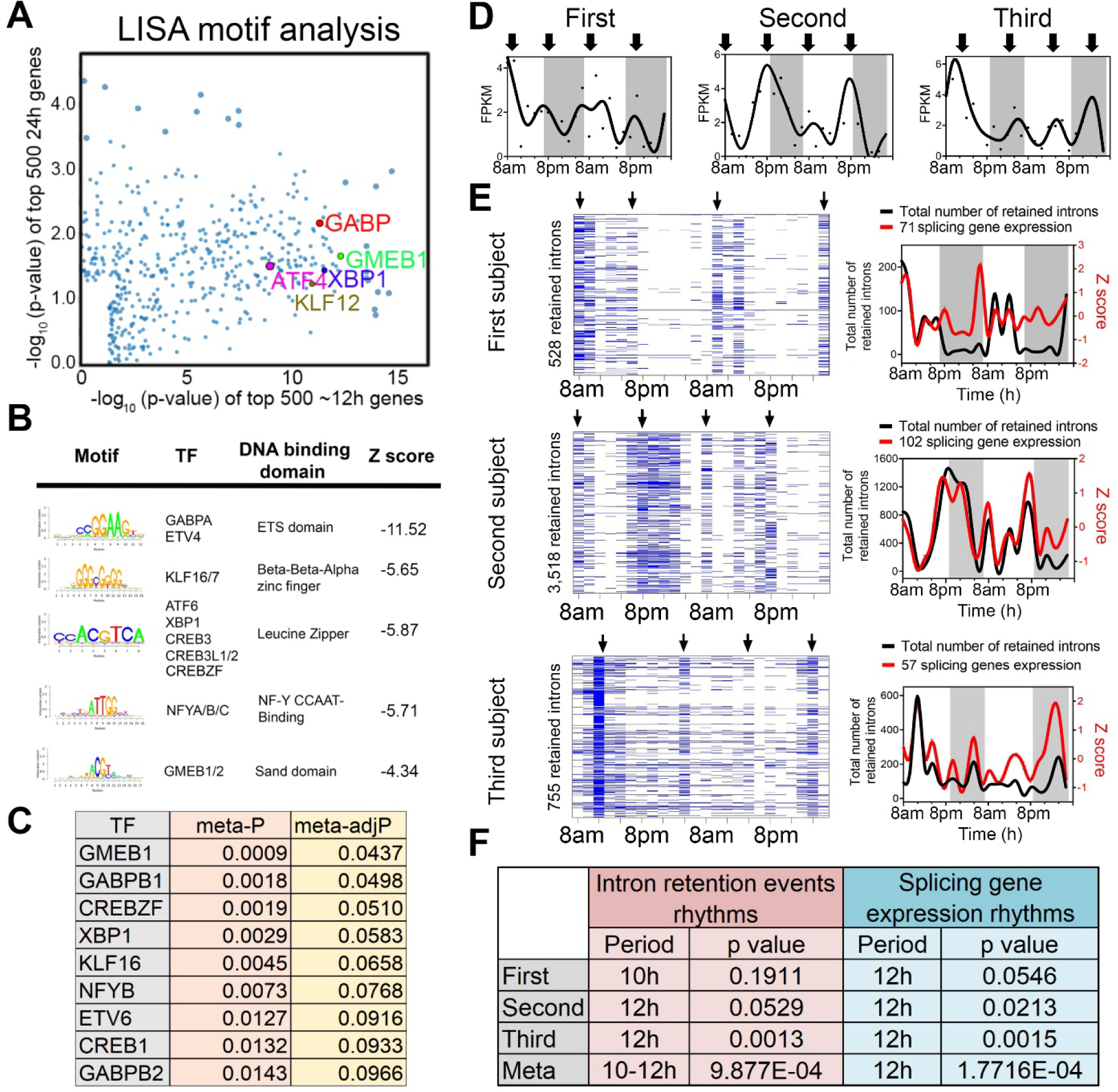
Regulatory and functional dissection of human ∼12h rhythms. (**A**) Scatter plot demonstrating the log normalized p values of motifs identified for top 500 (ranked by meta p values) ∼12h (x axis) versus the circadian genes using the LISA program, with selective TFs highlighted. (**B**) Top motifs enriched at the promoters of ∼top 500 12h genes using the SeqPos motif tool in Galaxy/Cistrome. (**C**) A table listing the TFs whose motifs are enriched in the promoters of ∼12h genes and whose gene expression also exhibit ∼12h rhythms with meta adj-P less than 0.1. (**D**) Temporal expression of *XBP1* in all three individuals. (**E, F**) Criterions for IR are set as T>=20, J>=1, FPKM>=2 and NE score>=0.9. Heatmap (left) and quantification (right) of temporal intron retention events, superimposed with the Z score normalized temporal expression of splicing genes exhibiting ∼12h rhythms (**E**). Statistics for IR and spicing gene expression ∼12h rhythms detection by RAIN (**F**).

We next incorporated a recently published XBP1 ChIPmentation dataset from mouse T helper cells into our analysis [40]. We found that XBP1 target genes—those genes directly bound by XBP1 and whose expression is reduced in the absence of XBP1—were significantly enriched in the putative ∼12h human gene set, but not in the circadian gene set (**Fig. S8A-C**). This included the mRNA processing genes *Tceb1*, *Thoc6*, *Smu1*, *Snrnp53*, *Eny2* and *Snrpc*, mitochondria metabolism-related genes such as *Gabpa*, *Mrpl40* and *Mrpl27,* and proteostasis genes *Mydgf*, *Rnf7*, *Serp1* as well as *Xbp1* itself (**Fig. S8A**). These results suggest XBP1, GABP and KLF transcription factor family members as candidate transcriptional regulators of ∼12h rhythms in human white blood cells, with XBP1 a strong candidate given its previously identified role as a major regulator of 12h transcriptional oscillations in the murine liver [13].

### ∼12h rhythms are synchronized to RNA splicing functionality

Given representation of RNA metabolism and mRNA splicing pathways in ∼12h transcriptional rhythms, we tested whether such oscillations would translate into a downstream functional effect in the form of alterations in mRNA splicing. We interrogated the RNA-seq data for evidence of rhythmicity in intron retention (IR) events, predicting that if the rhythm of RNA splicing genes’ transcription was functionally relevant, it would temporally correlate with global IR. We applied a recently published algorithm iREAD [41]. Using two different criteria for defining retained introns, we identified ∼12h rhythms of global IR events in all three participants (with meta-p<0.003) (**Figs. 3E**, **F and S9A-D**). IR rhythms were synchronized to the expression of mRNA splicing genes (**Figs. 3E**, **F and S9A-D**). These data suggest synchronization of mRNA splicing gene programs to splicing functionality, further implicating 12h rhythms in the regulation of RNA metabolism.

We next performed GO analysis on the set of transcripts in which we detected retained introns. While the GO terms associated with intron retention genes were largely consistent at the morning and evening peaks, we also observed differential enrichment in a subset. The morning intron retention gene sets were enriched in immune functions, especially those involving the display of intracellular peptide fragments to cytotoxic T cells via MHC class 1 complexes (*e.g.,* HLA-A/C/E and B2M), whereas the evening intron retention gene sets exhibited greater heterogeneity across the three participants (**Figs. S10, 11**). Taken together, these data suggest that one function of human ∼12h rhythms in peripheral leukocytes might be the anticipatory splicing of antigen presentation genes.

### Evolutionary conservation of ∼12h gene programs and function

∼12h rhythms in coastal and estuarine animals suggest an ancient evolutionary origin [6, 39, 42–46] (**Fig. 4A**). To test for homology between the hitherto identified human ∼12h rhythm genes and circatidal genes in marine animals, we compared the most robust ∼12h human genes (meta-adjP less than 0.10) with the circatidal gene program uncovered in *Aiptasia diaphana* [47], a sea anemone species which shares a eumetazoan ancestor with *Homo sapiens* ∼700 million years ago in the Cryogenian period [48]. In line with our human data, the most enriched biological pathways amongst *A. diaphana* circatidal genes were mRNA processing and protein metabolism, distinct from those enriched in the circadian genes, which included pathways of carbohydrate metabolism and detoxification [44] (**Fig. S12A**). We found 404 ∼12h genes common to both species, a number significantly higher than the expected value of 263 if there was no evolutionary conservation (p=4.6e-23 by Chi-square test) (**Fig. 4B**). This subset was also enriched in mRNA splicing and proteostasis pathways (**Fig. 4C-D**). We confirmed the evolutionary conservation of ∼12h rhythms of gene expression between human and *A. diaphana* by using two additional gene sets of human ∼12h genes: 653 gene set with meta-adjP values less than 0.05 (**Fig. 1C**), or the 851 gene set shared in all three individuals depicted in **Fig. S4B** (**Fig. S12B, C**). Protein processing in the ER was the most consistently enriched GO term using the different selection criteria for ∼12h genes (**Fig. S12B, C**). In contrast to ∼12h ultradian rhythms, we found modest evolutionary conservation of circadian gene expression between human (with meta-adjP value less than 0.10) and *A. diaphana* (p=0.049 by Chi-square test), with the 102 common circadian genes moderately enriched in carbohydrate metabolism and detoxification pathways (**Fig. S12D-F**).

**Fig. 4.**
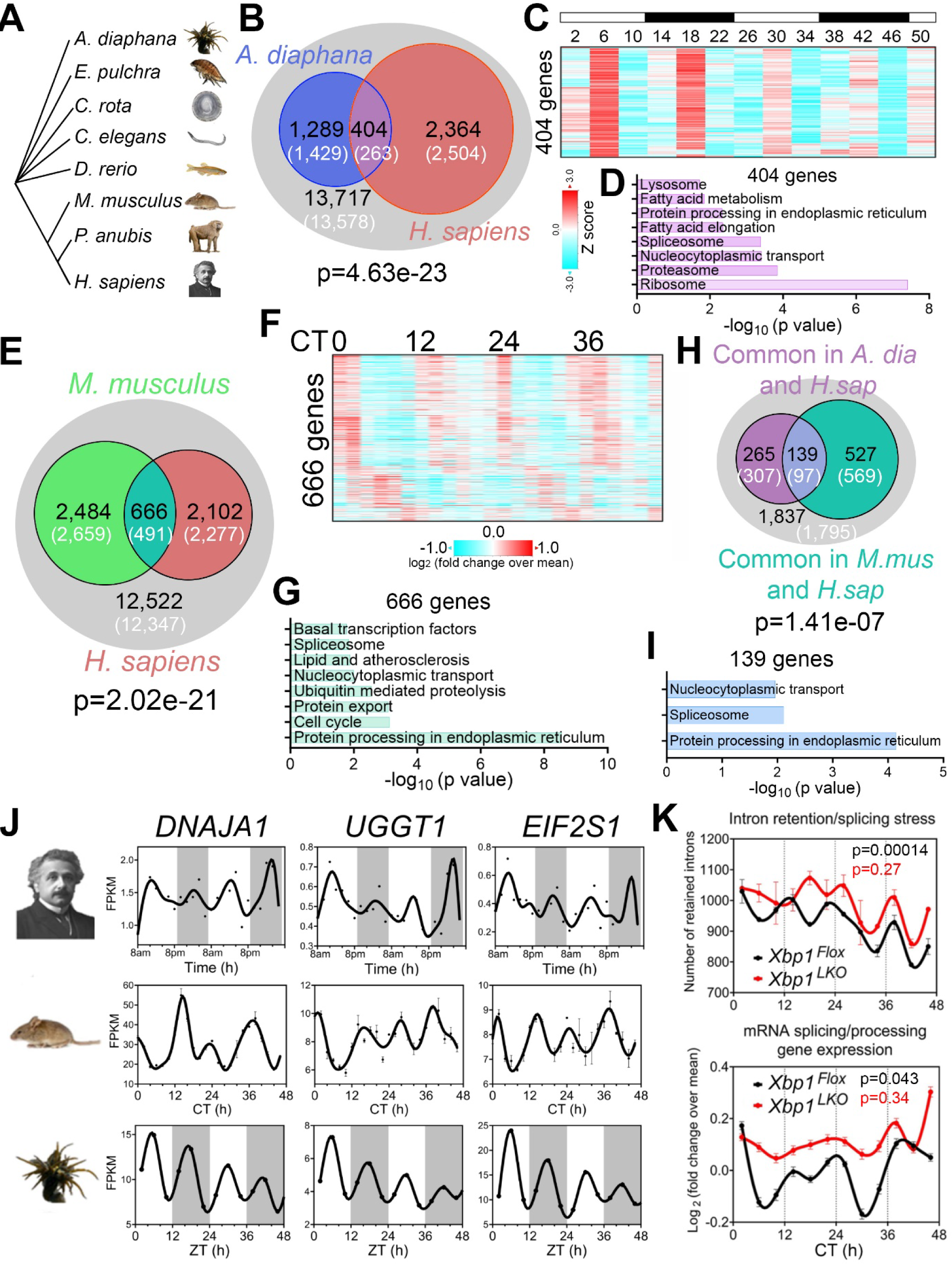
Evolutionary conservation of ∼12h gene programs. (**A**) Phylogenetic tree of select species for which ∼12h rhythms of gene expression have been demonstrated. *A. diaphana* is the most distant from *H. sapiens*. (**B**) Venn diagram comparing distinct and shared ∼12h genes in human (meta adj-P<0.1) and *A. diaphana* (reported in [44]). Only genes expressed in human white blood cells (denoted by the grey circle) are included in the analysis. Both observed and predicted number of genes (under the null hypothesis that ∼12h genes are not evolutionarily conserved and thus independently detected in these two species) are further shown. P value of 4.6e-23 is calculated by Chi-square test. (**C, D**) Heatmap of temporal expression (Z score normalized) of 404 circatidal genes in *A. diaphana* (**C**) and GO analysis of the top enriched pathways (**D**). (**E**) Venn diagram comparing distinct and shared ∼12h genes in human (meta adj-P<0.1) and mouse liver (reported in [13]). Only genes that are expressed in human white blood cells (denoted by the grey circle) are included in the analysis. Both observed and predicted number of genes (under the null hypothesis that ∼12h genes are not evolutionarily conserved and thus independently detected in these two species) are further shown. P value of 2.0e-21 is calculated by Chi-square test. (**F, G**) Heatmap of temporal expression (log 2 normalized) of 666 ∼12h genes in mouse liver (**F**) and GO analysis of the top enriched pathways (**G**). (**H**) Venn diagram further comparing the overlap of commonly identified 404 ∼12h genes in human and A. diaphana, and 666 ∼12h genes in human and mouse. Both observed and predicted number of genes (under the null hypothesis that ∼12h genes are not evolutionarily conserved and thus independently detected in all three species) are further shown. P value of 1.4e-7 is calculated by Chi-square test. (**I**) GO analysis of the top enriched pathways of 139 ∼12h genes commonly identified in all three species. (**J**) Representative temporal expression of evolutionarily conserved ∼12h genes in human, mouse, and *A. diaphana*. (**K**) ∼12h rhythms of intron retention events (top) and RNA splicing genes expression (bottom) are attenuated by liver specific loss of function of XBP1 (*Xbp1^LKO^*: red), relative to control (*Xbp1^Flox^*: black) in mice. P values calculated by RAIN for 12h rhythms.

If the 12-hour gene program is conserved from coastal invertebrates to humans, we reasoned that lower mammals like mice should also exhibit oscillations in similar gene programs. We re-examined a previously published temporal hepatic RNA-seq data set in mouse liver and found 666 ∼12h genes common to both human and mouse (p=2.02e-21 by Chi-square test), again highly enriched in mRNA splicing and protein processing pathways (**Fig. 4E-G**). We identified 139 ∼12h genes common to all three species: human, mouse, and *A. diaphana*, which exceeded the 97 expected by chance (p=1.41e-7 by Chi-square test) (**Fig. 4H**). These 139 genes are enriched in protein processing in ER, spliceosome, and nucleocytoplasmic transport, the three pivotal steps in central dogma information flow. Specific examples include DNAJA1, a heat shock protein 70 cochaperones [49], UGGT1 that serves as the predominant ER glycoprotein quality control sensor [50] and *EIF2S1*, encoding the alpha subunit of the translation initiation factor eIF2 protein complex [51] (**Fig. 4I-J**). These results are congruent with our previous work in murine liver in identifying a regulatory function for ∼12h oscillator in central dogma information flow [13].

Finally, we tested for synchronization between the ∼12h rhythms and RNA splicing in mice by performing intron retention analysis of our previously published temporal RNA-seq data from mouse liver [13]. Similar to the human data, we observed strong alignment between ∼12h rhythms in splicing gene expression and global intron retention events peaking at the two ‘rush hours’ (CT2 and CT14) (**Fig. 4K**). In addition, liver-specific genetic ablation of XBP1 (the transcriptional regulator of murine hepatic 12h oscillations) abolished ∼12h rhythms of splicing gene expression and intron retention rhythms, leading to constant intron retention across the day (**Fig. 4K**). These collective data support evolutionary conservation of a ∼12h gene program in humans related to RNA and protein metabolism.

## Discussion

In this study, we discovered a ∼12h regulatory network in healthy humans, controlling gene programs related to fundamental processes of mRNA and protein metabolism. Remarkably, conservation of ∼12h rhythms in humans extended to specific gene orthologs and functional pathways present in marine animals. Our discovery of one such 12h pathway, ‘mRNA splicing,’ provided an opportunity to establish functional significance through identification of corresponding 12h rhythms in global intron retention events. Moreover, pathway analyses of rhythmic intron retention suggests that one functional consequence of 12h rhythms in human white blood cells could be rhythmic antigen presentation by splicing of MHC class I gene transcripts. We further demonstrate conservation of a potential major 12h transcriptional regulator, *XBP1,* which regulates 12h gene oscillations in mice [13]. *XBP1* not only exhibited 12h rhythmicity in our human participants but was also implicated by gene regulatory network inferences as a lead candidate regulator of human 12h gene programs. What adaptive advantage might 12h rhythms of gene expression confer? We reason that the ancient ∼12h rhythms that evolved in marine species in response to tidal cues have been coopted by humans as an adaptive response to accommodate physiological transitions related to feeding, physical activity, and sleep that are temporally concentrated at dawn and dusk.

While our study provides evidence of 12h rhythms of gene expression in humans inclusive of potential transcriptional regulators, it does not establish causality, nor does it fully address their relationship with the circadian clock. At least three mutually non-exclusive mechanisms have been proposed to explain the origin and regulatory mechanisms of ∼12h rhythms in mice, namely that they are not cell-autonomous and controlled by a combination of the circadian clock and environmental cues [25, 52, 53], that they are regulated by two anti-phase circadian transcriptional factors in a cell-autonomous manner [54], or that they are established by a cell-autonomous ∼12-hour oscillator [12, 13, 26, 28, 55, 56]. Future studies will be required to test whether candidate 12h regulatory factors are causal drivers of 12h gene expression rhythms and any direct relationship with the circadian clock. Such mechanistic studies are challenging in humans because prospective genetic manipulations cannot be performed. However, several potential future directions hold promise to advance a mechanistic basis for 12h rhythms, including longitudinal profiling of DNA binding by putative 12h transcriptional regulators at genome scale, temporal transcriptomic/cistromic profiling of human cohorts that exhibit genetic variation in candidate 12h regulatory factors, and assessment of the modifying effects of combinatorial genetic manipulation of circadian clock and candidate 12h regulators in human cells/tissues *ex vivo*.

Similar studies of molecular rhythms in model organisms typically require terminal tissue collection and therefore require inclusion of several animals at each timepoint [12, 13]. Given the genetic and environmental variability of free-living humans, analogous human studies have required hundreds of participants [57]. Our study strength owes to the prospective, repeated sample collection at high temporal resolution, which enabled high fidelity testing for a ∼12h gene program *within* each participant. Despite this strength, our study also demonstrates the challenge of identifying specific rhythmic genes, as several canonical circadian genes did not meet statistical significance in all participants. Even genes that are under rhythmic control may have additional confounding regulatory inputs from environmental or behavioral stimuli and as such we propose that functional relevance is enhanced by our pathway level analyses, which demonstrated commonality of the putative ∼12h pathways across all three human participants and their conservation in organisms ranging from marine animals to humans. While these data provide evidence in support of a ∼12h gene program in humans, larger studies will be needed to determine how variable the ∼12h programs are in the human population, the degree to which they are sensitive to aging or environmental stressors as observed for circadian rhythms, and most importantly whether disruption of ∼12h rhythms is a causal determinant of disease pathobiology. Aside from the identification of ∼12h pathways related to fundamental cellular processes, several pieces of evidence suggest that ∼12h rhythms are important for maintenance of homeostasis: (i) when we adjusted for detection sensitivity by matching sampling frequency to period length, the number of putative ∼12h genes was similar to the circadian program; and (ii) prior human studies suggest ∼12h rhythms in physiological metrics of relevance to human health and disease, including heart rate variability, blood pressure, hormone levels, and cognitive function [6, 8, 58, 59]. As such, our discovery of ancient ∼12h gene programs in humans provides rationale for future studies to determine whether ∼12h rhythms should be viewed alongside circadian rhythms as a core molecular determinant of health and disease.

## Materials and Methods

### Human participants and study protocol

The study protocol was approved by the University of Pittsburgh Institutional Review Board (Study 20020034; approval date: 6/4/2020) and written consent was obtained from all study participants. We studied 3 healthy male participants, who were recruited through online advertisements. Inclusion criteria consisted of individuals 18-35 years of age with a self-reported regular nighttime sleep schedule and a body mass index between 18.5-24.9 kg/m^2^. Volunteers were excluded if they admitted to nighttime shift work or other regular nighttime sleep-disrupting activities, if they had any chronic medical conditions, took any medications or recreational drugs, or used tobacco products. Potential volunteers presented for a screening visit, inclusive of measurement of body weight, height, BMI, and laboratory studies, including a comprehensive metabolic panel (electrolytes, kidney function and liver function tests), complete blood count, 25-OH vitamin D, and thyroid stimulating hormone level, to screen for potential subclinical chronic diseases. We excluded participants with low hemoglobin/hematocrit, abnormal thyroid function and individuals with 25-OH vitamin D < 20 ng/mL.

Qualifying study participants maintained a food diary, which was used to estimate their daily caloric intake and subsequently presented to the University of Pittsburgh Medical Center (UPMC) Clinical Translational Research Center (CTRC) for a 3-day inpatient visit. On the morning of admission, participants selected items from a food menu designed to match their standard daily caloric intake with a uniform macronutrient composition of 55% carbohydrates, 25% fat, 20% protein per day. No interventions were performed during the first 24-hour period of acclimatization.

On the morning of the second day at 8am, an intravenous (IV) line was placed. Blood samples were then collected every two hours for 48 hours (total 24 samples). Nighttime blood collection was performed through a long IV line from outside the room so that the study participant would not be exposed to light or woken up during blood collection. Blood was then immediately processed in the Center for Human Integrative Physiology, two floors above the UPMC CTRC by a rotating study team, all of whom were trained in the processing procedures for this study. Blood was centrifuged (1900 RCF x 10 min) and the buffy coats were collected and immediately snap frozen in liquid nitrogen for storage at -80C.

### RNA-Seq and data analysis

RNA was isolated from peripheral blood buffy coat samples on the automated Chemagic 360 (Perkin Elmer) instrument according to the manufacturer’s instructions. Extracted RNA was quantitated by Qubit™ RNA BR Assay Kit (Thermo Fisher Scientific) followed by the RNA quality check using Fragment Analyzer (Agilent). For each sample, RNA libraries were prepared from 100ng RNA using the KAPA RNA HyperPrep Kit with RiboErase (Kapa Biosystems) according to manufacturer’s protocol, followed by quality check using Fragment Analyzer (Agilent) and quantification by qPCR (Kapa qPCR quantification kit, Kapa biosystem) on the LightCycler 480 (Roche). The libraries were normalized and pooled, and then sequenced using NovaSeq6000 platform (Illumina) to an average of 40M 101bp paired end reads per sample. Low-quality reads and adapter sequences were trimmed from the raw sequencing data with Trimmomatic [60]. The remaining reads were then aligned to human reference genome hg38 with STAR aligner [61]. Gene counts were quantified with the STAR-quantMode GeneCounts function. Fragments per kilobase of transcript per million mapped fragments (FPKM) were quantified with Cufflinks [62].

### Identification of the oscillating transcriptome

Averaged FPKM values at each time were used for cycling transcripts identification. Lowly expressed transcripts were removed by calculating the background expression in each participant using the average expression of a panel of 62 genes known not to be expressed in peripheral blood cells (**Table S2**). Temporal transcriptomes were participant to linear detrend prior to identification of oscillations by either the eigenvalue/pencil or RAIN methods. For the eigenvalue/pencil method [10, 12], a maximum of four oscillations were identified for each gene. Criterion for circadian genes were: period between 20h to 25h for first and second participants and 24h to 30h for the third participant, decay rate between 0.8 and 1.2; for ∼12h genes: period between 9.6h to 13.6h for the second and third participants and 10h to 14h for the first participant, decay rate between 0.8 and 1.2; for ∼8h genes: period between 6h to 8h for the first participant and 7h to 9h for the second participant, decay rate between 0.8 and 1.2; for ∼16h genes; period between 14h to 18h for the third participant. The relative amplitude was calculated by dividing the amplitude by the mean expression value for each gene. To determine FDR, we used a permutation-based method that randomly shuffles the time label of gene expression data and subjected each permutation dataset to the eigenvalue/pencil method applied with the same criterion [29]. These permutation tests were run 5,000 times, and FDR was estimated by taking the ratio between the mean number of rhythmic profiles identified in the permutated samples (false positives) and the number of rhythmic profiles identified in the original data. Analyses were performed in MatlabR2017A. RAIN analysis was performed as previously described in Bioconductor (3.4) (http://www.bioconductor.org/packages/release/bioc/html/rain.html) with either 48h continuous data or 24h data with biological duplicates as input [11]. We included genes exhibiting a period between 10h and 14h with a p value less than 0.05 as having ∼12h expression in all three participants. FDR was calculated using the Benjamini-Hochberg procedure. Heat maps were generated with Gene Cluster 3.0 and TreeView 3.0 alpha 3.0 using Z score normalized values.

For meta-analysis, we used Fisher’s method, which combines extreme value probabilities from each test, commonly known as “p-values”, into one test statistic (X2) using the formula

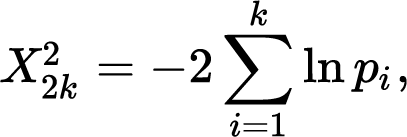

where pi is the p-value for the ith hypothesis test.

For RAIN analysis on temporal IR events and splicing gene expression, raw data was subjected to polynomial detrend (n=2) before RAIN analysis.

### Defining oscillating genes

The eigenvalue method can detect multiple superimposed oscillations. Therefore, we defined a circadian gene as one that exhibited a circadian rhythm, regardless of its amplitude relative to other superimposed oscillations. Similar criteria were applied to other oscillations. As such, a gene can meet criteria for both a circadian and ∼12h gene. By comparison, we define a dominant circadian gene as one in which the superimposed circadian rhythm has the largest amplitude among all oscillations. With this definition, dominant circadian and dominant 12h genes are mutually exclusive.

### Intron retention detection

Intron retention events were detected by tool iREAD [41]. Intron retention events are selected either with default settings T>=20, J>=1, FPKM>=2 and NE score>=0.9 or more stringent settings where T>=20, J>=1, FPKM>=3 and NE score>=0.9.

### Gene ontology analysis

DAVID (Version 2021) [63] (https://david.ncifcrf.gov) was used to perform Gene Ontology analyses. Briefly, gene names were first converted to DAVID-recognizable IDs using Gene Accession Conversion Tool. The updated gene list was then subject to GO analysis using all Homo Sapiens as background and with Functional Annotation Chart function. GO_BP_DIRECT, KEGG_PATHWAY or UP_KW_BIOLOGICAL_PROCESS was used as GO categories. Only GO terms with a p value less than 0.05 were included for further analysis.

### Motif analysis

Motif analysis was performed with the SeqPos motif tool (version 0.590) embedded in Galaxy Cistrome using all motifs within the homo sapiens reference genome hg19 as background. LISA analysis was performed using webtool (http://lisa.cistrome.org/).

## Acknowledgments

We thank Toren Finkel for critical discussion and insightful advice on the manuscript, and Soukaina Eljamri for help with the human study protocols. This research was supported in part by the University of Pittsburgh Center for Research Computing through the resources provided. Specifically, this work used the HTC cluster, which is supported by NIH award number S10OD028483.

## Author contributions

Conceptualization: BZ, PKF, MLS

Methodology: BZ, SL, PKF, MLS

Investigation: BZ, SL, PKF, MLS, NLD, NKD, WD, LBS, TA, REA, GVNK, SI, TP, AXS, MS

Visualization: BZ, SL Formal Analysis: BZ, SL, HL

Funding acquisition: BZ, PKF, MLS

Resources: BZ, PKF, MLS

Software: BZ, SL

Validation: BZ, SL

Supervision: BZ, PKF, MLS

Writing – original draft: BZ, SL, PKF, MLS

Writing – review & editing: all authors

## Data and materials availability

All raw and processed sequencing data generated in this study have been submitted to the NCBI Gene Expression Omnibus (GEO; http://www.ncbi.nlm.nih.gov/geo/) under accession numbers GSE220120.

## Supplementary Materials for

**Fig. S1.**
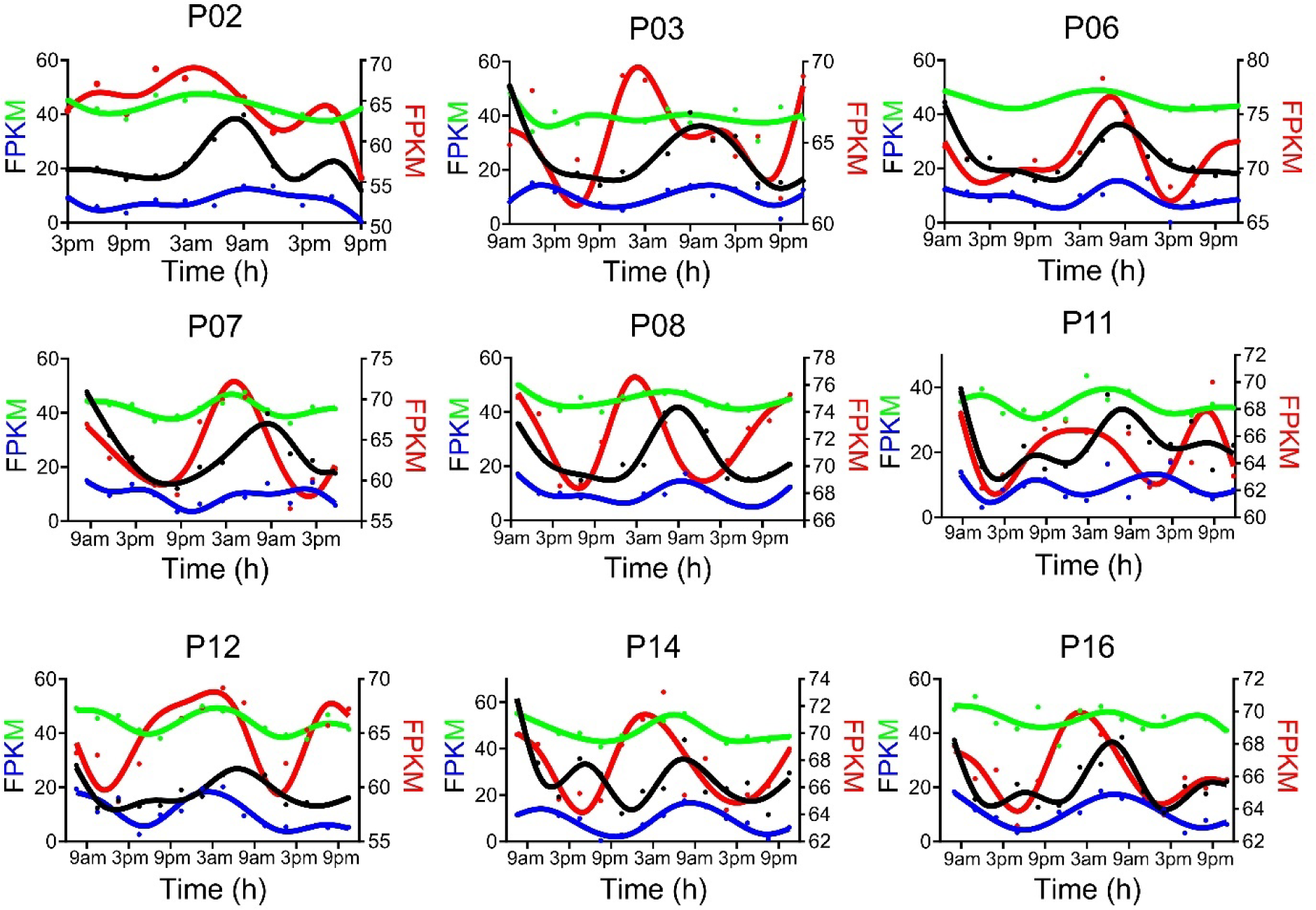
Inter-individual variability of circadian rhythms of gene expression reported in the Wittenbrink et al JCI study. Raw temporal expression profile (dot) and spline fit (solid line) of *PER1* (black), *PER2* (blue), *TSPAN4* (green) and *LGALS3* (red) genes in nine different individuals, reported in [14].

**Fig. S2.**
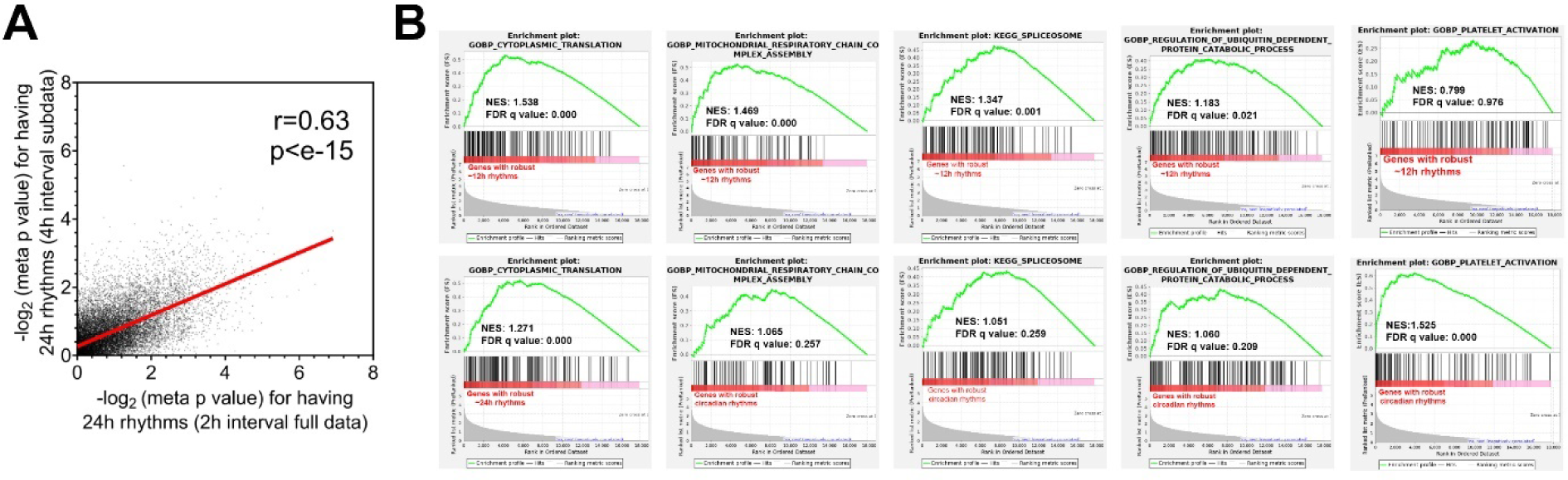
Genes with ∼12h and circadian rhythms are enriched in distinct biological pathways. (**A**) Scatter plot comparing log normalized meta p values for having circadian rhythms for each gene when the full 2h sampling interval dataset (x-axis) or the 4h sampling interval subset (y-axis) was used for the analysis. (**B**) GSEA showing enrichment scores for different gene sets in ∼12h (top) and circadian (bottom) genes.

**Fig. S3.**
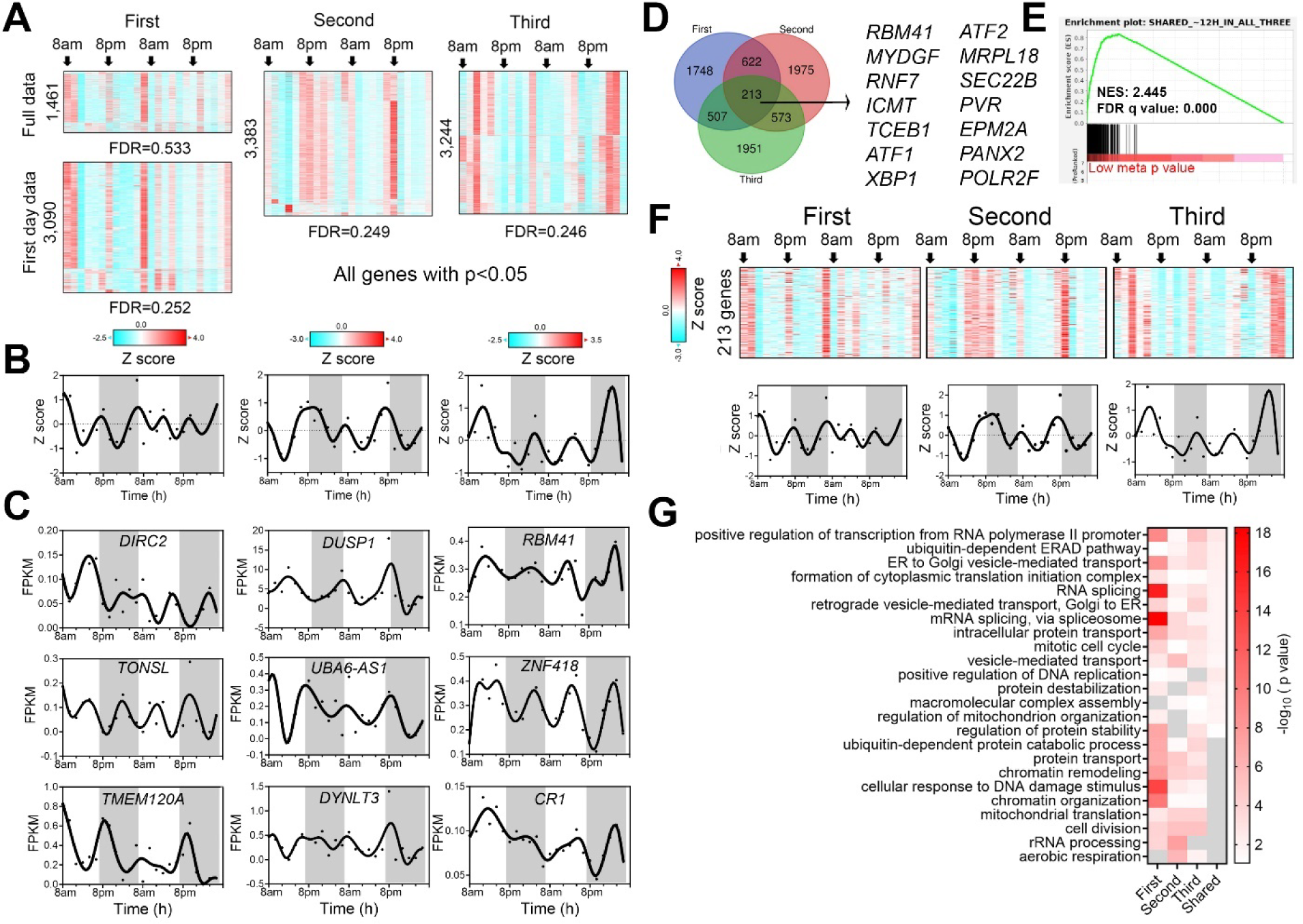
RAIN analyses of 48h temporal transcriptome to detect ∼12h genes. A single data point was inputted for each sample collected over the 48h protocol. (**A**) Heatmap of ∼12h genes from all three participants with respective p values smaller than 0.05. The identification of a greater number of ∼12h genes with lower FDR in the first participant with restriction of inputted data to the first 24 hours indicates the ∼12h rhythm is dampened in the second day in this participant, consistent with what was found with the eigenvalue/pencil method. (**B**) Quantification of the average expression (Z score normalized) of genes shown in A, with raw data (dots) and spline fit (solid lines) shown. (**C**) Raw temporal expression (dot) profile and spline fit (solid line) of top three ∼12h genes with the smallest p values in each of the three individuals. (**D**) Venn diagram depicting common and distinct ∼12h genes for each individual, with selective common genes shown on the right. (**E**) GSEA showing enrichment score of 213 common ∼12h genes on robust ∼12h genes ranked by meta p values. (**F**) Heatmap and quantification of 213 common ∼12h genes uncovered from all three participants. (**G**) GO analysis of all and shared ∼12h genes in each individual.

**Fig. S4.**
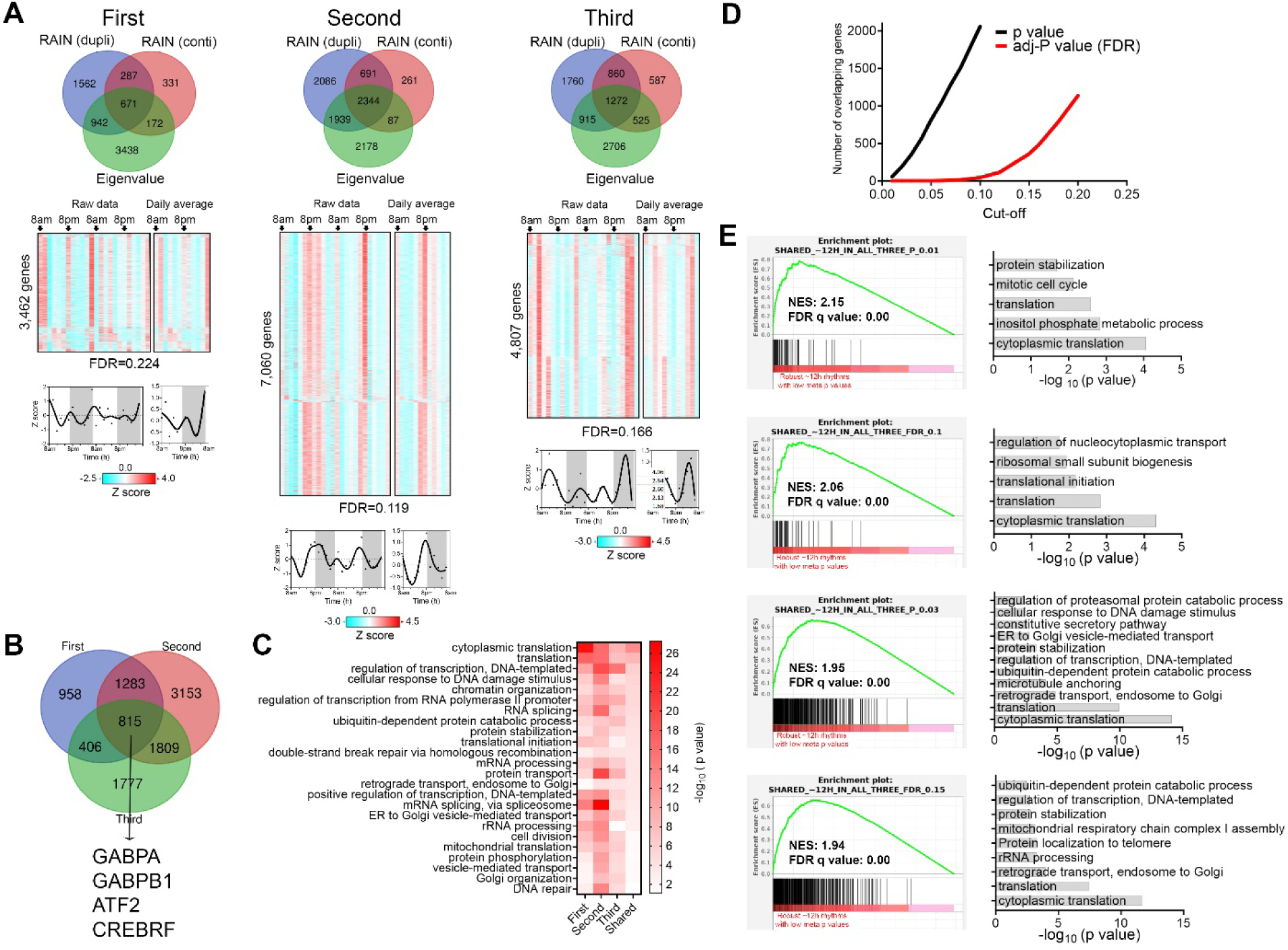
RAIN analyses of temporal transcriptomes using consecutive 24 hours datasets as biological replicates. (**A**) Venn diagram comparing the ∼12h programs uncovered by the three methods (top) and heatmap and quantification of ∼12h genes uncovered by the RAIN dupli method in all three participants (bottom). (**B**) Venn diagram depicting common and distinct ∼12h genes uncovered in each participant. (**C**) GO analysis of all and shared ∼12h genes in all three participants. (**D**) Scatter plot comparing the number of overlapping ∼12h genes between the three individuals against different p values or FDR cut-offs for all individuals. (**E**) GSEA showing enrichment scores for four gene sets of overlapping ∼12h genes with p<0.01 (56 genes), p<0.03 (358 genes), FDR<0.1 (46 genes) and FDR<0.15 (361 genes) cut-off (left) on ∼12h genes ranked by meta-p values and GO analysis of these genes (right).

**Fig. S5.**
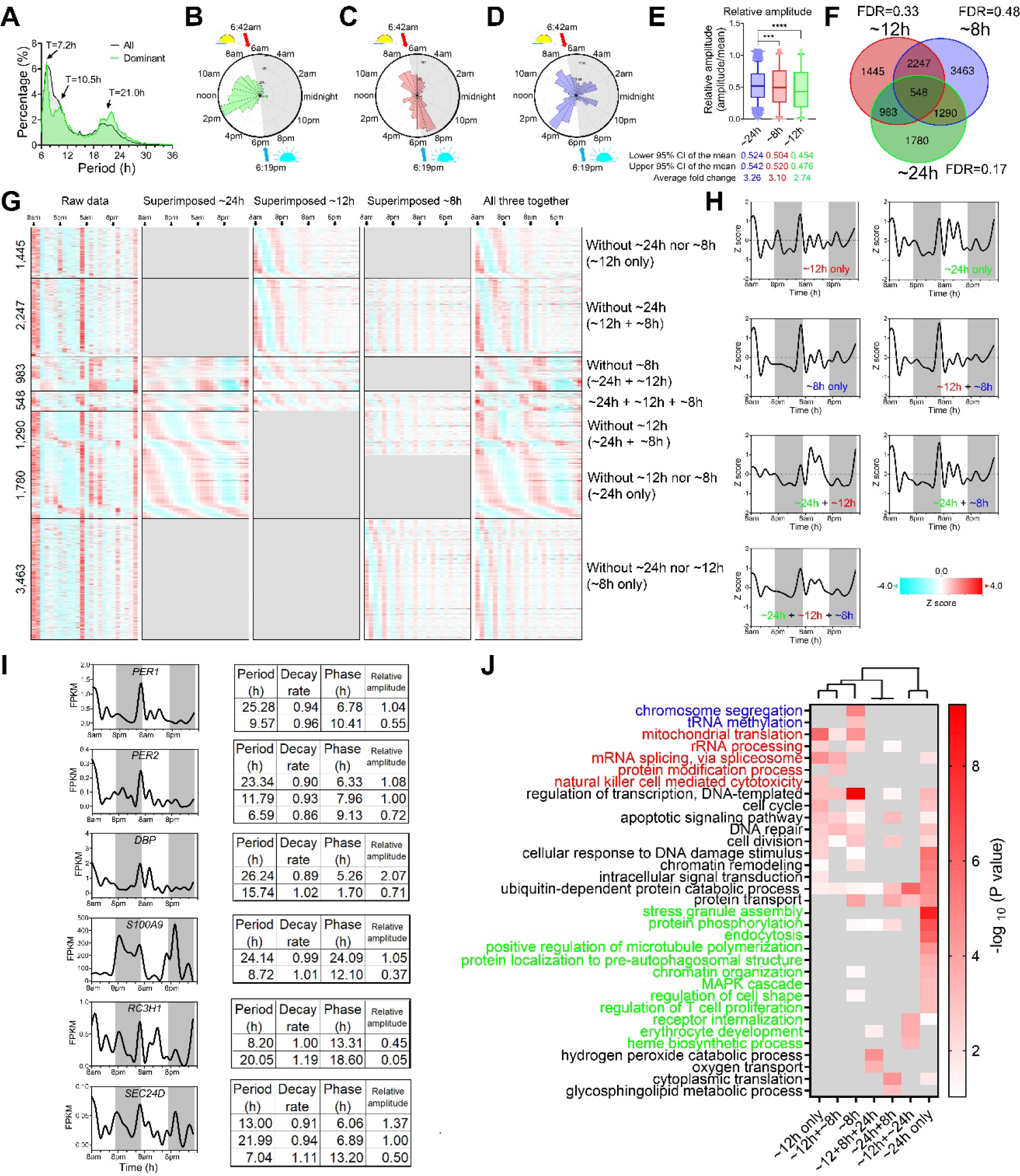
Transcriptome spectrum of first human participant. (**A**) Period distribution of all and dominant oscillations. (**B-D**) Polar histogram illustrating the phase distribution of ∼24h (**B**), ∼12h (**C**) and ∼8h (**D**) oscillations. (**E**) Relative amplitude (mean-normalized) of ∼8h, ∼12h and ∼24h oscillations. (**F**) Venn diagram showing distinct and shared oscillations for each period. (**G, H**) Heatmap (**G**) and quantification (**H**) of decomposition of raw temporal transcriptome into harmonics cycling at ∼8h, ∼12h or ∼24h periods. (**I**) Representative temporal expression of select genes and eigenvalue/pencil decomposition uncovering all superimposed oscillations for each gene. (**J**) GO analysis of all genes in **G**.

**Fig. S6.**
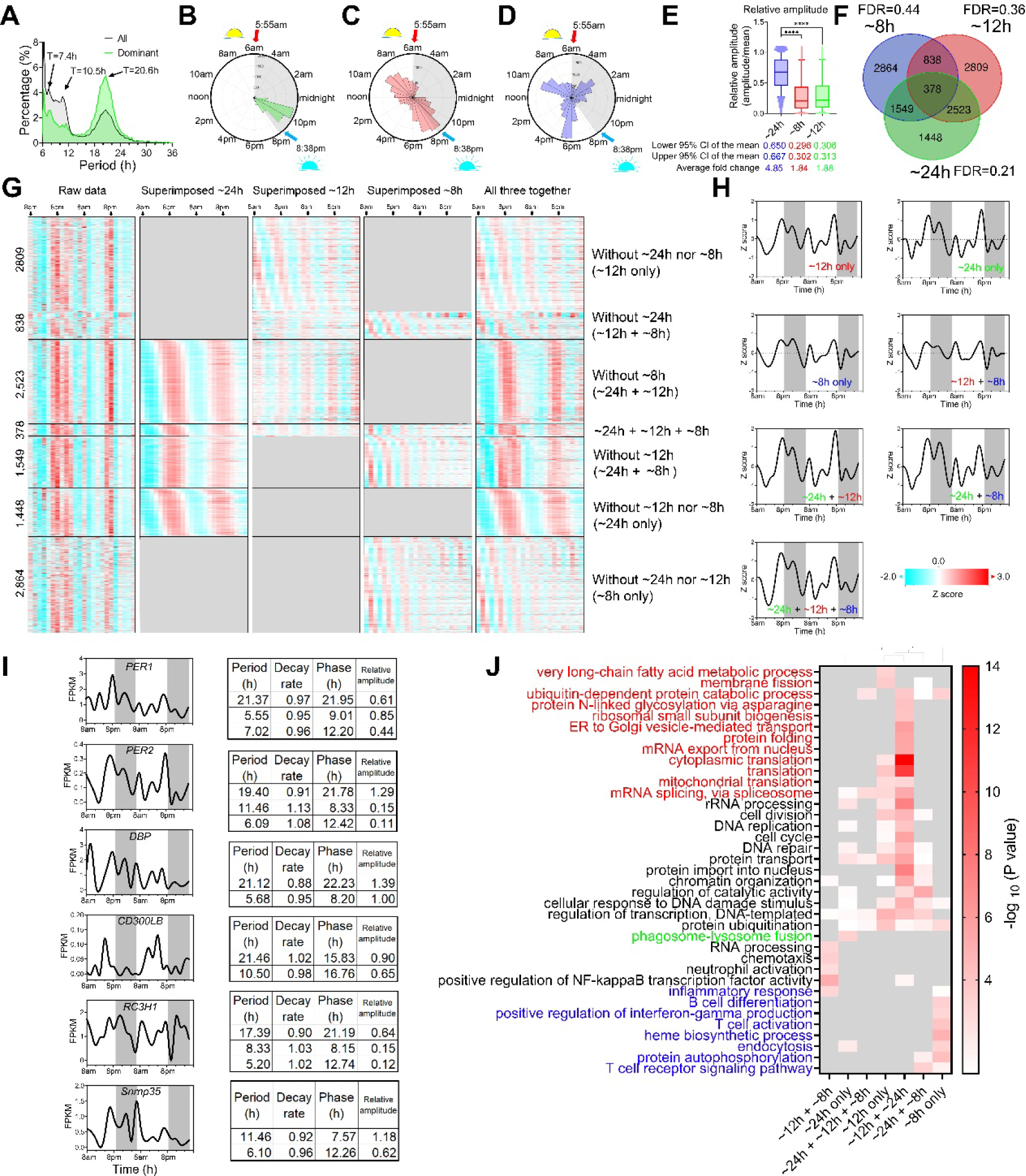
Transcriptome spectrum of the second human participant. (**A**) Period distribution of all and dominant oscillations. (**B-D**) Polar histogram illustrating the phase distribution of ∼24h (**B**), ∼12h (**C**) and ∼8h (**D**) oscillations. (**E**) Relative amplitude (mean-normalized) of ∼8h, ∼12h and ∼24h oscillations. (**F**) Venn diagram showing distinct and shared oscillations for each period. (**G, H**) Heatmap (**G**) and quantification (**H**) of decomposition of raw temporal transcriptome into harmonics cycling at ∼8h, ∼12h or ∼24h periods. (**I**) Representative temporal expression of selective genes and eigenvalue/pencil decomposition uncovering all superimposed oscillations for each gene. (**J**) GO analysis of all genes in **G**.

**Fig. S7.**
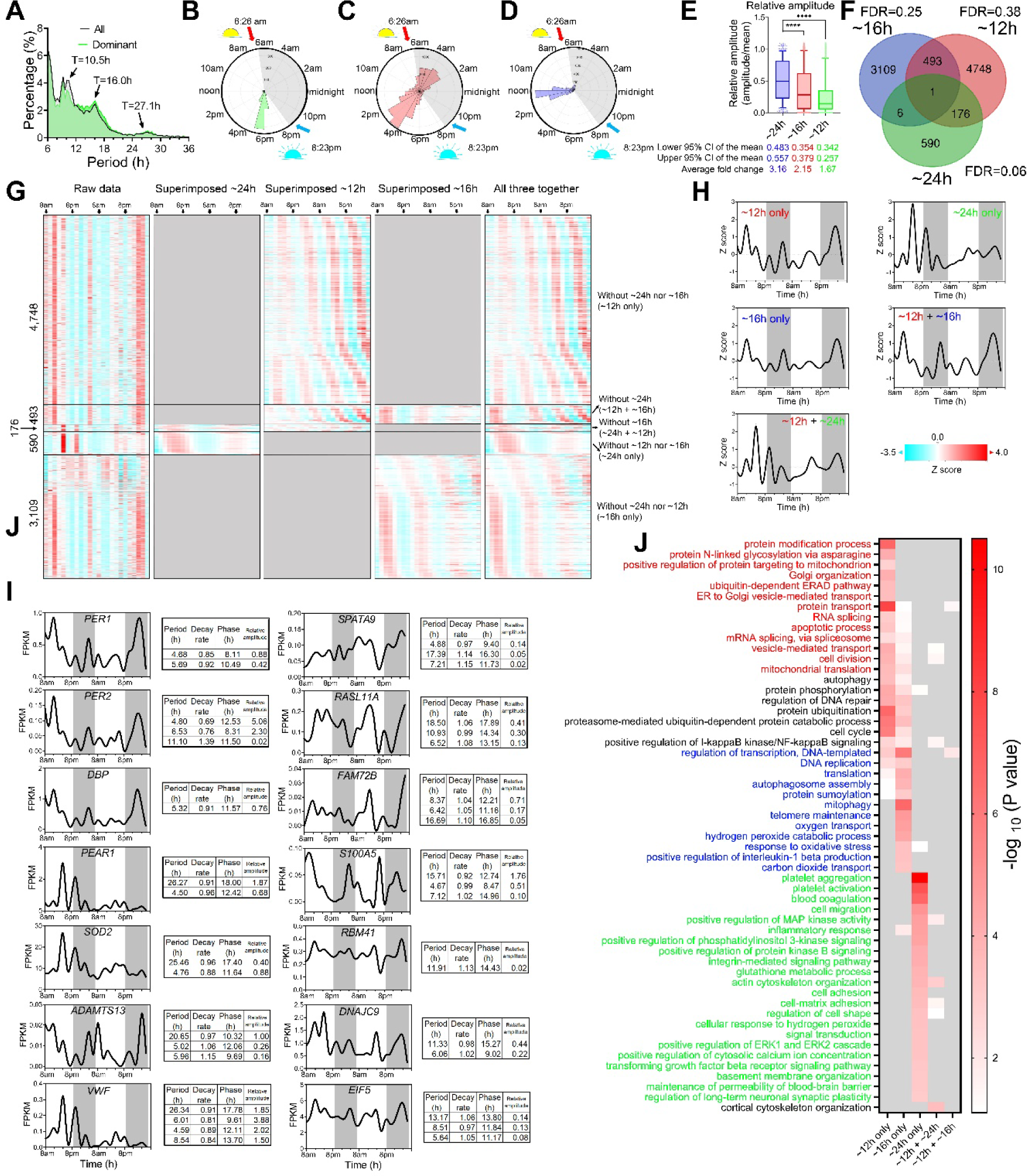
Transcriptome spectrum of the third human participant. (**A**) Period distribution of all and dominant oscillations. (**B-D**) Polar histogram illustrating the phase distribution of ∼24h (**B**), ∼12h (**C**) and ∼16h (**D**) oscillations. (**E**) Relative amplitude (mean-normalized) of ∼16h, ∼12h and ∼24h oscillations. (**F**) Venn diagram showing distinct and shared oscillations for each period. (**G, H**) Heatmap (**G**) and quantification (**H**) of decomposition of raw temporal transcriptome into oscillations cycling at ∼16h, ∼12h or ∼24h periods. (**I**) Representative temporal expression of selective genes and eigenvalue/pencil decomposition uncovering all superimposed oscillations for each gene. (**J**) GO analysis of all genes in **G**.

**Fig. S8.**
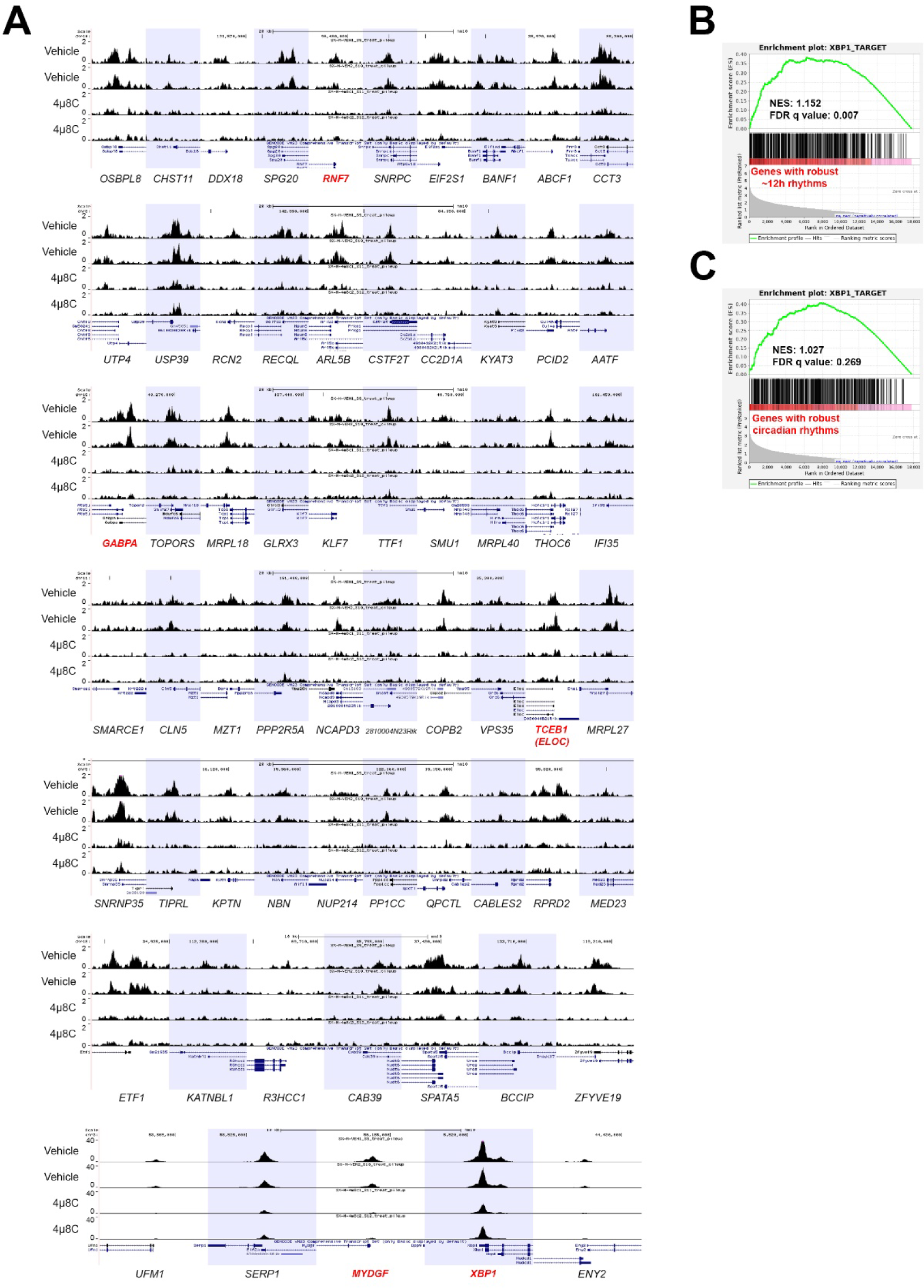
XBP1 chromatin binding landscape of 12h genes in mouse Th2 cells. (**A**) UCSC genome browser view of XBP1 chromatin recruitment to the promoters of a selected set of ∼12h genes in murine Th2 cells. (**B, C**) GSEA showing enrichment score of top 500 XBP1 target genes in Th2 cells (ranked by fold reduction of target gene expression with XBP1 deletion) in robust ∼12h (**B**) or circadian (**C**) genes identified in humans.

**Fig. S9.**
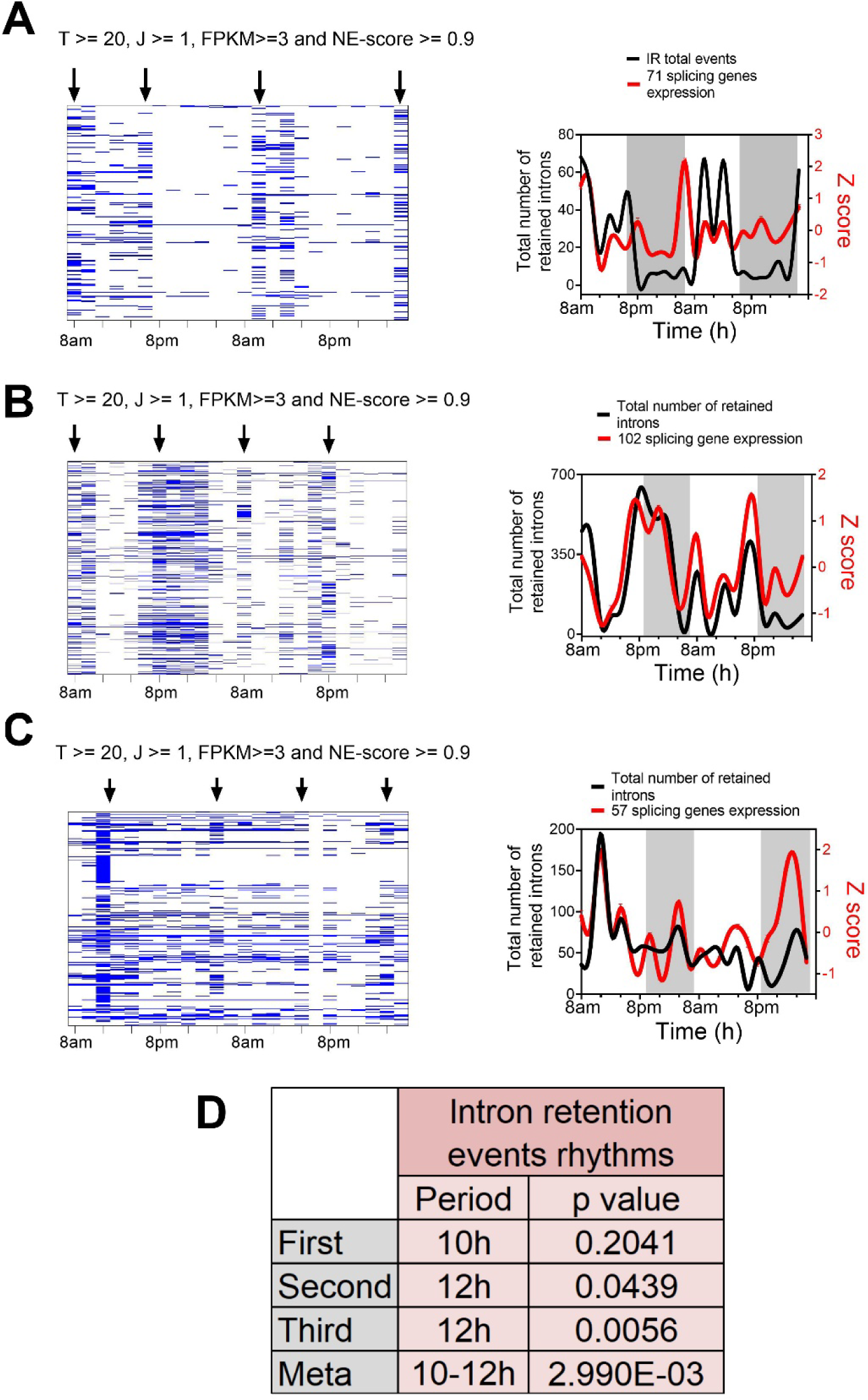
∼12h rhythms are synchronized to global intron retention rhythms. Criterions for IR are set as T>=20, J>=1, FPKM>=3 and NE score>=0.9. (**A-C**) Heatmap (left) and quantification (right) of temporal IR events, superimposed with the Z score normalized temporal expression of ∼12h splicing gene expression in the first (**A**), second (**B**) and third (**C**) participants. IR events are detected with the more stringent setting of the iRead algorithm as described in the methods section. (**D**) Statistics for IR ∼12h rhythms detection by RAIN.

**Fig. S10.**
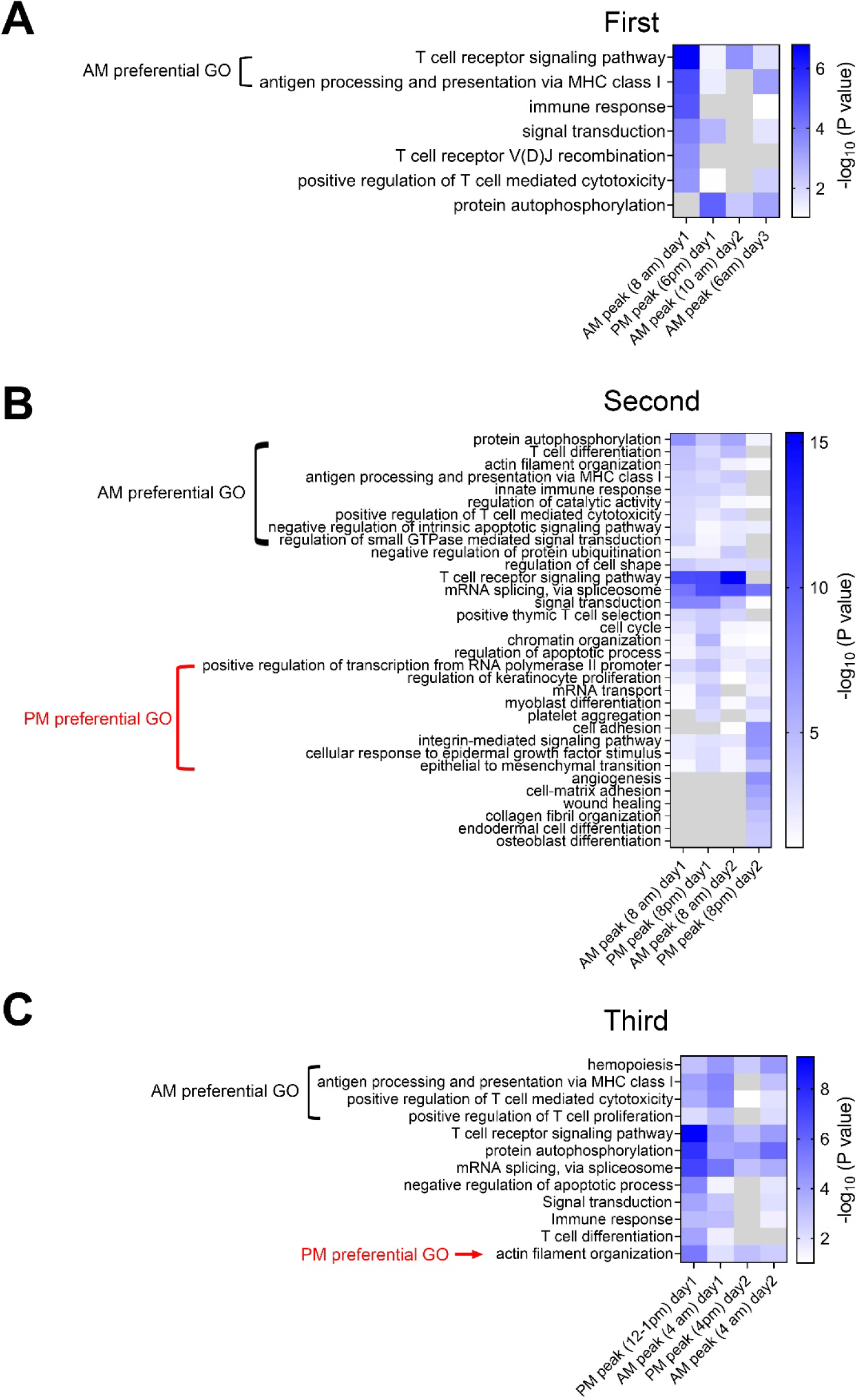
Genes with intron retentions are enriched in immune genes. Intron retention events are selected with default settings T>=20, J>=1, FPKM>=2 and NE score>=0.9. GO analysis of IR genes at different peaks in the first (**A**), second (**B**) and third (**C**) participant.

**Fig. S11.**
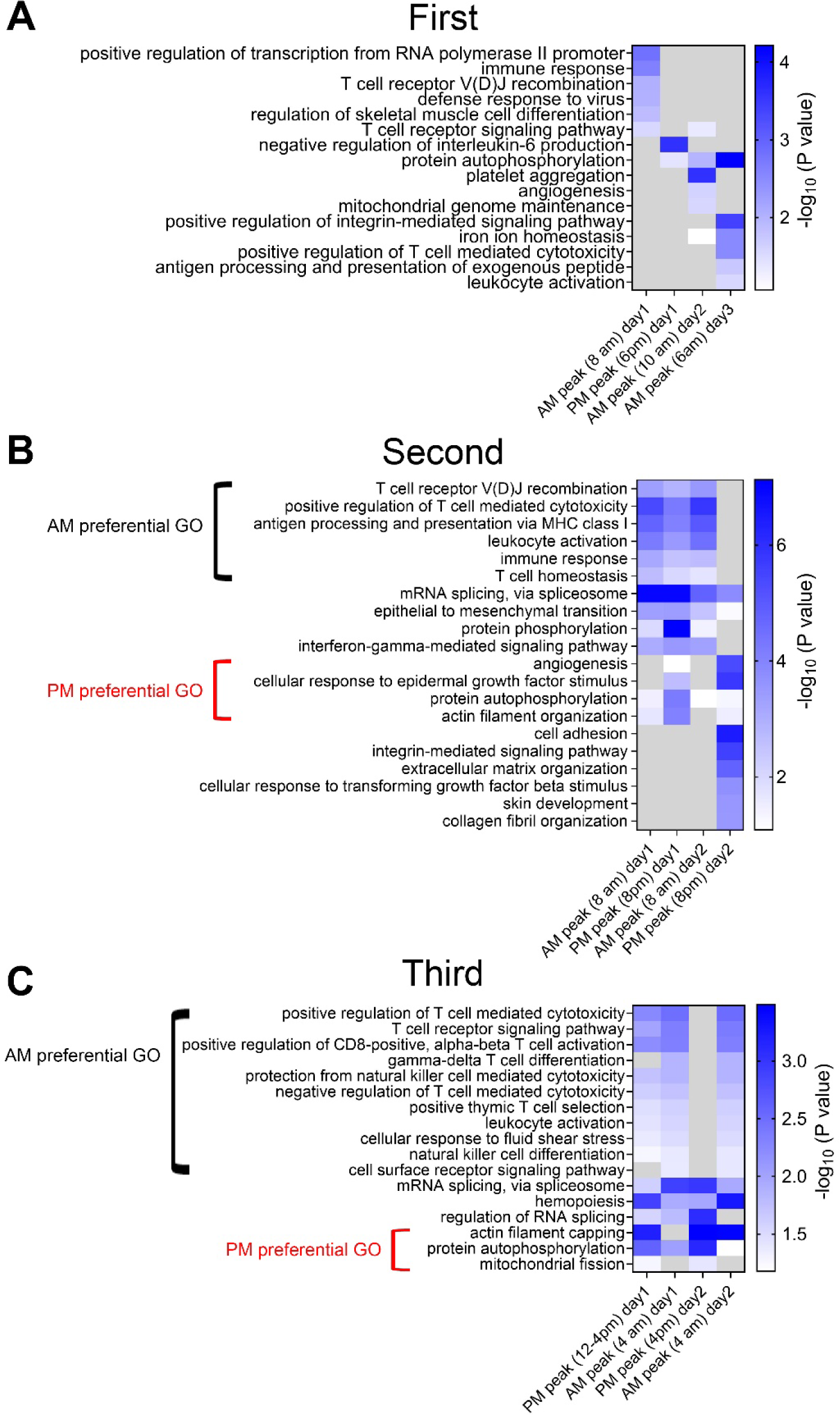
Genes with intron retentions (more stringent criterion) are enriched in immune genes. Intron retention events are selected with default settings T>=20, J>=1, FPKM>=3 and NE score>=0.9. GO analysis of IR genes at different peaks in the first (**A**), second (**B**) and third (**C**) participant.

**Fig. S12.**
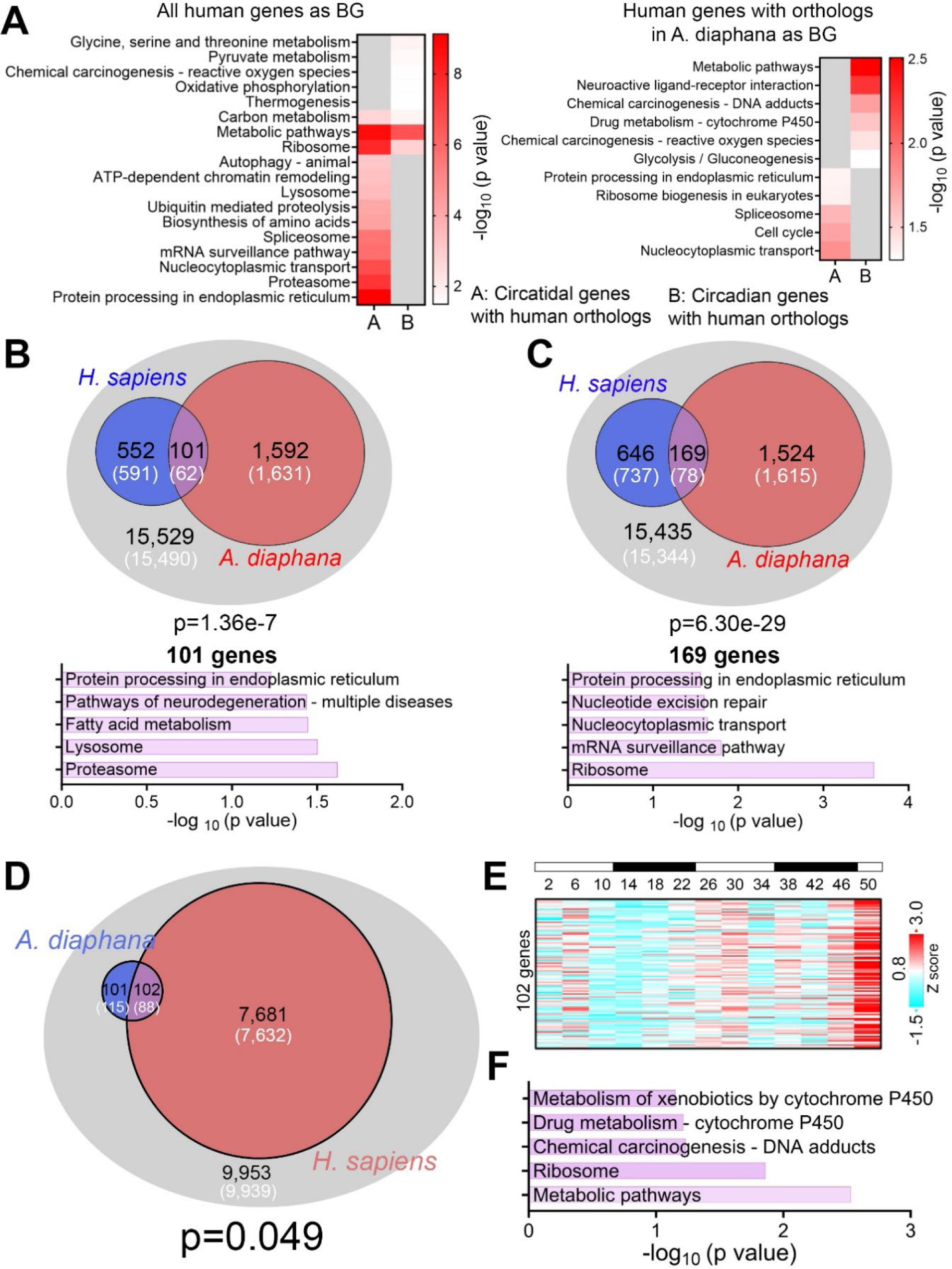
Evolutionary conservation of ∼12h, but not circadian gene programs between A. diaphana and human. (**A**) GO analysis of all circatidal genes in A. diaphana that have human orthologs using all human genes (left) or only those with A. diaphana orthologs (right) as background. (**B, C**) Venn diagram comparing distinct and shared ∼12h genes in human (653 12h genes with meta adj-P<0.05 in **B** and 851 ∼12h genes reported in Fig. S4B in **C** and *A. diaphana* (reported in [44]). Only genes that are expressed in human white blood cells (denoted by the grey circle) are included in the analysis. Both observed and predicted number of genes (under the null hypothesis that ∼12h genes are not evolutionarily conserved and thus independently detected in these two species) are further shown. P values are calculated by the Chi-square test. GO analysis showing enriched KEGG terms for the 101 and 169 common genes, respectively. (**D**) Venn diagram comparing distinct and shared circadian genes in human (meta adj-P<0.1) and *A. diaphana* (reported in [44]). Only genes that are expressed in human white blood cells (denoted by the grey circle) are included in the analysis. Both observed and predicted number of genes (under the null hypothesis that circadian genes are not evolutionarily conserved and thus independently detected in these two species) are further shown. P value of 0.049 is calculated by Chi-square test. (**E, F**) Heatmap of temporal expression (Z score normalized) of 102 circadian genes in *A. diaphana* (**E**) and GO analysis of the top enriched pathways (**F**).

Table S1. RPKM quantification of temporal expression for the human participants

Table S2. RAIN results for ∼12h genes and meta-analysis of circadian genes.

Table S3. Eigenvalue/pencil results

